# UBAP2L-dependent coupling of PLK1 localization and stability during mitosis

**DOI:** 10.1101/2022.10.24.513562

**Authors:** Lucile Guerber, Aurore Vuidel, Yongrong Liao, Charlotte Kleiss, Erwan Grandgirard, Izabela Sumara, Evanthia Pangou

**Author notes:** **Correspondence:** Evanthia Pangou and Izabela Sumara. Institute of Genetics and Molecular and Cellular Biology (IGBMC), Illkirch, France. These authors contributed equally to this work (Lucile Guerber, Evanthia Pangou).

## Abstract

PLK1 is an important regulator of mitosis whose protein levels and activity fluctuate during the cell cycle. PLK1 dynamically localizes to various mitotic structures to regulate chromosome segregation. However, the signaling pathways linking localized PLK1 activity to its protein stability remain elusive. Here, we identify the Ubiquitin-Binding Protein 2-Like (UBAP2L) that controls both, the localization and the protein stability of PLK1.We demonstrate that UBAP2L is a spindle-associated protein whose depletion leads to severe mitotic defects. UBAP2L depleted cells are characterized by increased PLK1 protein levels and abnormal PLK1 accumulation in several mitotic structures such as kinetochores, centrosomes and mitotic spindle. UBAP2L deficient cells exit mitosis and enter the next interphase in the presence of aberrant PLK1 kinase activity. The C-terminal domain of UBAP2L mediates its function on PLK1 independently of its role in stress response signaling. Importantly, the mitotic defects of UBAP2L depleted cells are largely rescued upon chemical inhibition of PLK1. Overall, our data suggest that UBAP2L is required to finetune the ubiquitin-mediated PLK1 turnover during mitosis as a means to maintain genome fidelity.

## Introduction

Protein kinases represent key regulatory elements of the mitotic cycle, transferring phosphorylation signals to critical effectors (Nigg, 2001). Polo-like kinase 1 (PLK1) represents one of the key mitotic enzymes ensuring both mitotic entry as well as fidelity of genome segregation, mitotic exit and cytokinesis (Schmucker & Sumara, 2014; Combes *et al*, 2017; Petronczki *et al*, 2008). PLK1 is a serine/threonine kinase with an enzymatic domain at its N-terminal and a Polo-Box domain (PBD) at its C-terminal part. PDB is a unique feature of the PLK kinase family and confers specificity to phosphorylation substrates (Strebhardt, 2010; Barr *et al*, 2004; Zitouni *et al*, 2014). The expression of PLK1 is cell cycle dependent, with its levels peaking at G2/M transition and declining during mitotic exit and early G1 (Golsteyn *et al*, 1995; Bruinsma *et al*, 2012). At the end of mitosis, PLK1 is subjected to proteolytic ubiquitylation by the E3 ubiquitin ligase anaphase promoting complex/cyclosome (APC/C), which leads to its proteasomal degradation (Lindon & Pines, 2004).

PLK1 undergoes several post-translational modifications which finetune its dynamic localization, stability and activation/inactivation at several structures including centrosomes, kinetochores, central spindle and midbody (Schmucker & Sumara, 2014). During G2/M transition, PLK1 is enriched at centrosomes where it triggers pericentrin phosphorylation and the recruitment of pericentriolar material (PCM) components, thereby promoting centrosome maturation and separation (Lee & Rhee, 2011). From prometaphase until metaphase, PLK1 is enriched at kinetochores via interaction with phosphorylated kinetochore receptors including the mitotic checkpoint serine/threonine-protein kinase BUB1 beta (BubR1) and the inner centromere protein (INCENP) (Elowe *et al*, 2007; Goto *et al*, 2006; Qi *et al*, 2006). At kinetochores, PLK1 regulates stability of kinetochore-microtubule (KT-MT) attachments (Elowe *et al*, 2007) and spindle assembly checkpoint (SAC) function (Sumara *et al*, 2004; Lenart *et al*, 2007). During anaphase, PLK1 migrates to the spindle midzone through interaction and phosphorylation of PRC1, facilitating chromosome segregation (Hu *et al*, 2012). Finally, during telophase and cytokinesis PLK1 accumulates in the midbody, where it regulates ingression of cleavage furrow and separation of daughter cells (Petronczki *et al*, 2007).

Our previous studies have shown that PLK1 is a substrate for non-proteolytic ubiquitylation by the E3 ligase Cullin3 (CUL3) (Beck *et al*, 2013). CUL3 in complex with the substrate specific adaptor Kelch-like protein 22 (KLHL22) mono-ubiquitylates PLK1 within its PBD domain and interferes with phospho-receptors’ binding, leading to the timely removal of PLK1 from kinetochores prior to anaphase and faithful genome segregation. This modification is counteracted by the opposing function of the deubiquitylase (DUB) USP16 (Zhuo *et al*, 2015) that promotes proper chromosome alignment in early mitosis. Thus, both dynamic localization and protein stability of PLK1 are tightly regulated by phosphorylation- and ubiquitylation-based signals to ensure proper mitotic progression and genome stability. However, the exact molecular mechanisms linking the regulation of localized PLK1 activity to its protein stability, as well as which signaling pathways trigger relocalization of PLK1 from early to late mitotic structures remain poorly characterized.

Ubiquitin-Binding Protein 2-Like (UBAP2L, also known as NICE-4) is a highly conserved ubiquitin- and RNA-binding protein with versatile roles in multiple signaling cascades and cellular functions (Guerber *et al*, 2022). While UBAP2L has been mostly studied in the context of stress response signaling (Cirillo *et al*, 2020; Huang *et al*, 2020), recent evidence suggests that it can be involved in regulating mitotic progression (Maeda et al., 2016). UBAP2L is methylated within its RGG domain located at the N-terminal part and this modification was reported to promote the stability of KT-MT attachments, ensuring accurate chromosome distribution (Maeda et al., 2016). However, it remains unknown whether additional mechanisms to the reported methylation can actively drive the role of UBAP2L in cell division and what is the identity of direct downstream targets of UBAP2L during mitosis.

In this study we provide evidence that UBAP2L regulates both dynamic localization and protein stability of PLK1 independently of the RGG domain. We demonstrate that UBAP2L controls the global turnover of PLK1 throughout mitotic progression by controlling its proper localization at centrosomes, kinetochores and mitotic spindle. UBAP2L itself localizes at the mitotic spindle and centrosomes during early mitotic stages and accumulates in the midzone and midbody during anaphase and telophase respectively. Cells lacking UBAP2L are characterized by mitotic delay, aberrant chromosome segregation and micronuclei formation. Depletion of UBAP2L impairs the dynamic removal of individual PLK1 pools from different mitotic structures and reduces poly-ubiquitylation and degradation of PLK1. Consequently, depletion of UBAP2L leads to increased PLK1 stability after completion of mitosis, resulting in high PLK1 kinase activity in the subsequent interphase. Importantly, the mitotic defects of UBAP2L depleted cells are largely restored upon PLK1 inhibition, suggesting that the genomic instability observed in absence of UBAP2L can be directly coupled to aberrant PLK1 mitotic signaling.

## Results

### UBAP2L regulates proper chromosome segregation during mitosis

To identify novel ubiquitin-related factors with a potential role in mitosis, we previously performed a high-content visual siRNA screen in HeLa cells for human ubiquitin-binding domain (UBD) proteins (Krupina et al., 2016) and assessed the phenotypes of irregular nuclear shape that are often the result of chromosome segregation defects (Jevtić *et al*, 2014). UBAP2L was among the top hits of the screen (Krupina *et al*, 2016), as its deletion resulted in increased number of cells with polylobed nuclei and multinucleation, phenotypes highly comparable to those observed upon down-regulation of the positive control CUL3 (Fig EV1A) (Maerki *et al*, 2009; Sumara *et al*, 2007). Interestingly, UBAP2L has been proposed to be involved in mitotic progression via its methylation by the arginine methyltransferase PRMT1, which is required for KT-MT attachment formation and chromosome alignment (Maeda et al., 2016) but the direct downstream targets of UBAP2L that are important for mitotic progression are currently unknown. In order to confirm our screening results and to further dissect the precise role of UBAP2L during mitosis, we deleted UBAP2L in HeLa cells using CRISPR/Cas9-mediated gene editing (Fig EV1B and C) and performed time-lapse live video microscopy (Fig 1A and Movies EV1-3). UBAP2L Knock-Out (KO) cells displayed significant delay in mitotic onset and in prophase to anaphase time course compared to isogenic wild-type (WT) control cells (Fig 1A-C). In addition, UBAP2L KO cells were characterized by chromosome alignment defects and DNA bridges during anaphase and telophase, after which the cells either exited mitosis as multinucleated cells or died after prolonged mitotic arrest (Fig 1A, and 1D-F). All mitotic phenotypes observed upon UBAP2L loss were further confirmed upon acute depletion of UBAP2L by siRNA treatment (Fig1H-M and Movies EV4-6). Moreover, the presence of micronuclei and nuclear atypia in UBAP2L depleted cells was further confirmed in additional cell lines derived from colorectal cancer (DLD-1) (Fig EV1D and E) and osteosarcoma (U2OS) (Fig EV1F and G), respectively. Overall, our results suggest that UBAP2L regulates proper and timely chromosome segregation during mitosis.

**Figure 1.**
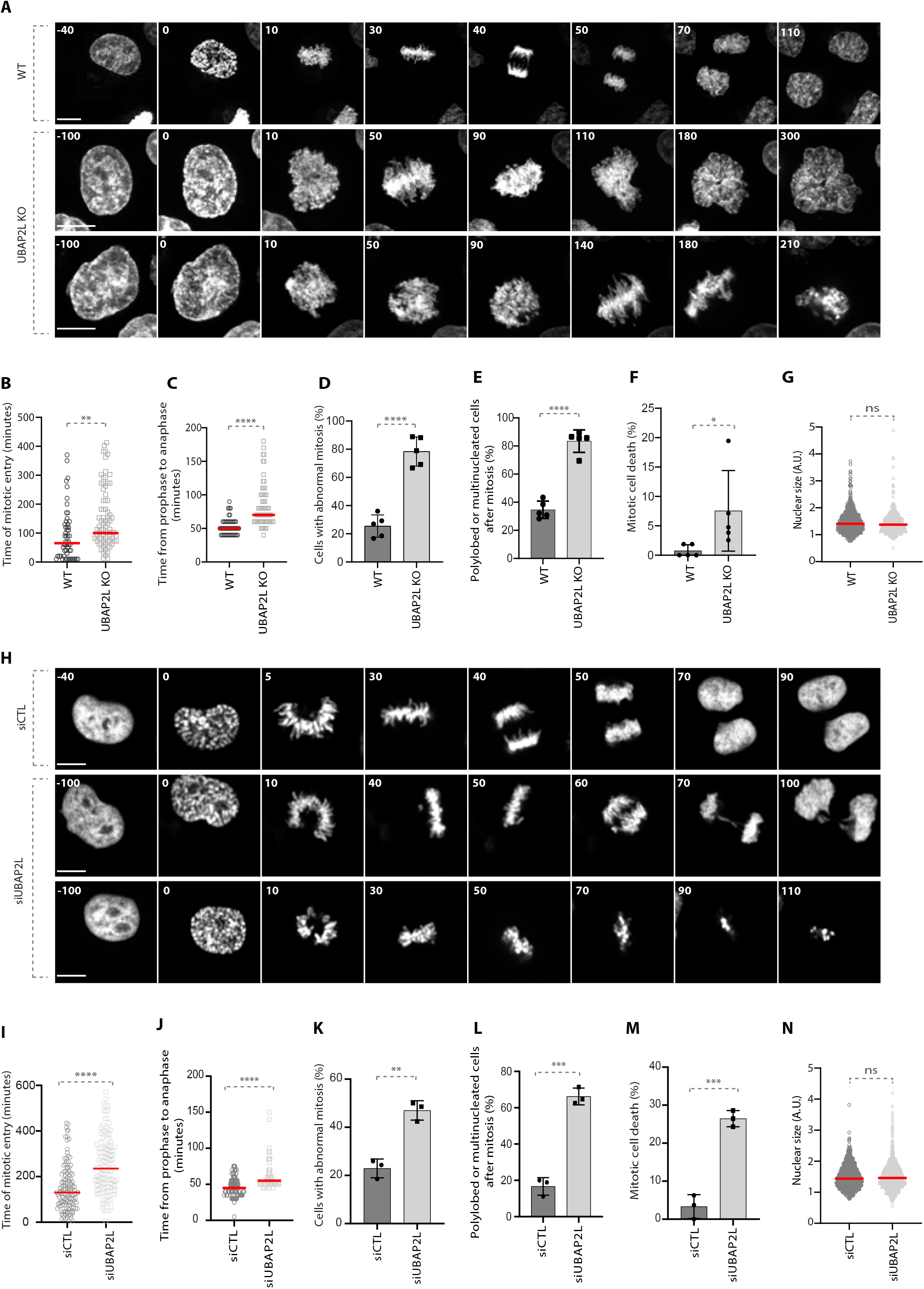
UBAP2L regulates proper chromosome segregation during mitosis. **A**. Spinning disk time-lapse microscopy of WT and UBAP2L KO HeLa cells synchronized with double thymidine block and release (DTBR) in mitosis. The selected frames of the movies are depicted and the corresponding time is indicated in minutes. SiR-DNA was used for DNA staining. Scale bars, 5 μm and 8μm. **B, C**. The time of mitotic entry **(B)** and from prophase to anaphase **(C)** was quantified. At least 50 cells per condition were analyzed for each experiment (n=5). Red bars represent the mean. Statistics were determined using the Mann-Whitney test (**P<0,01, ****P<0,0001). **D-F**. The percentages of cells with abnormal mitosis (chromosome misalignments and/or DNA bridges) **(D)**, polylobed or multinucleated daughter cells **(E)** and mitotic cell death **(F)** were quantified. At least 50 cells per condition were analyzed. Graphs represent the mean of five biological replicates ± standard deviation (SD) (two sample two-tailed t-test *P<0,05, ****P<0,0001). **G**. The nuclear size of at least 600 WT and UBAP2L KO interphasic cells was measured. Red bars represent the mean. Statistics were determined by the Mann-Whitney test (ns=non-significant). **H**. Spinning disk time-lapse microscopy of HeLa cells transfected with control (siCTL) or UBAP2L siRNA (siUBAP2L) for 48h and synchronized with DTBR in mitosis. The selected frames of the movies are depicted and the corresponding time is indicated in minutes. SiR-DNA was used for DNA staining. Scale bar, 6μm. **I, J**. The time of mitotic entry **(I)** and from prophase to anaphase **(J)** was quantified. At least 50 cells per condition were analyzed for each experiment (n=3). Red bars represent the mean. Statistics were determined using the Mann-Whitney test (****P<0,0001). **K-M**. The percentages of cells with abnormal mitosis (chromosome misalignments and/or DNA bridges) **(K)**, polylobed or multinucleated daughter cells **(L)** and mitotic cell death **(M)** were quantified. At least 50 cells per condition were analyzed. Graphs represent the mean of three biological replicates ± SD (two sample two-tailed t-test **P<0,01, ***P<0,001). **N**. The nuclear size of at least 1000 control and UBAP2L-downregulated interphasic cells was measured. Red bars represent the mean. Statistics were determined by the Mann-Whitney test (ns=non-significant).

To exclude that the phenotypes observed in UBAP2L deficient cells are the secondary consequences of defects in a previous mitosis, we examined their ploidy status. Polyploidy was assessed by analyzing the nuclear size in WT versus UBAP2L KO cells (Fig 1G) and in cells treated with control siRNA (siCTL) or siRNA against UBAP2L (siUBAP2L) (Fig 1N). No significant differences in nuclear size were observed in either case, suggesting that mitotic defects observed in UBAP2L depleted cells are not caused by prior defective mitoses.

### UBAP2L localizes at the mitotic spindle but it does not regulate the spindle formation

Since chromosome alignment defects are often accompanied by aberrant centrosome numbers (Gönczy, 2015) and are known to induce spindle positioning defects (Tame *et al*, 2016), we wondered whether loss of UBAP2L affects the number of centrosomes and/or the polarity of the mitotic spindle. To this end, we assessed the centrosome number in WT and UBAP2L KO cells (Fig 2A-C), as well as in cells treated with siCTL or siUBAP2L (Fig 2D-F), using pericentrin as a centrosome marker. Our results on cells synchronized in G1/S phase (Fig 2A, B, D and E) and/or in early mitosis (prophase) (Fig 2A, C, D and F) showed that UBAP2L depletion does not lead to aberrant centrosome numbers. Finally, as revealed by α-tubulin staining, UBAP2L KO cells were able to form a bipolar mitotic spindle during metaphase despite the presence of segregation errors (Fig 2G, H). Our results indicate that the mitotic defects observed upon UBAP2L depletion are likely not the consequence of preexisting centrosome amplification and that UBAP2L does not regulate spindle formation.

**Figure 2.**
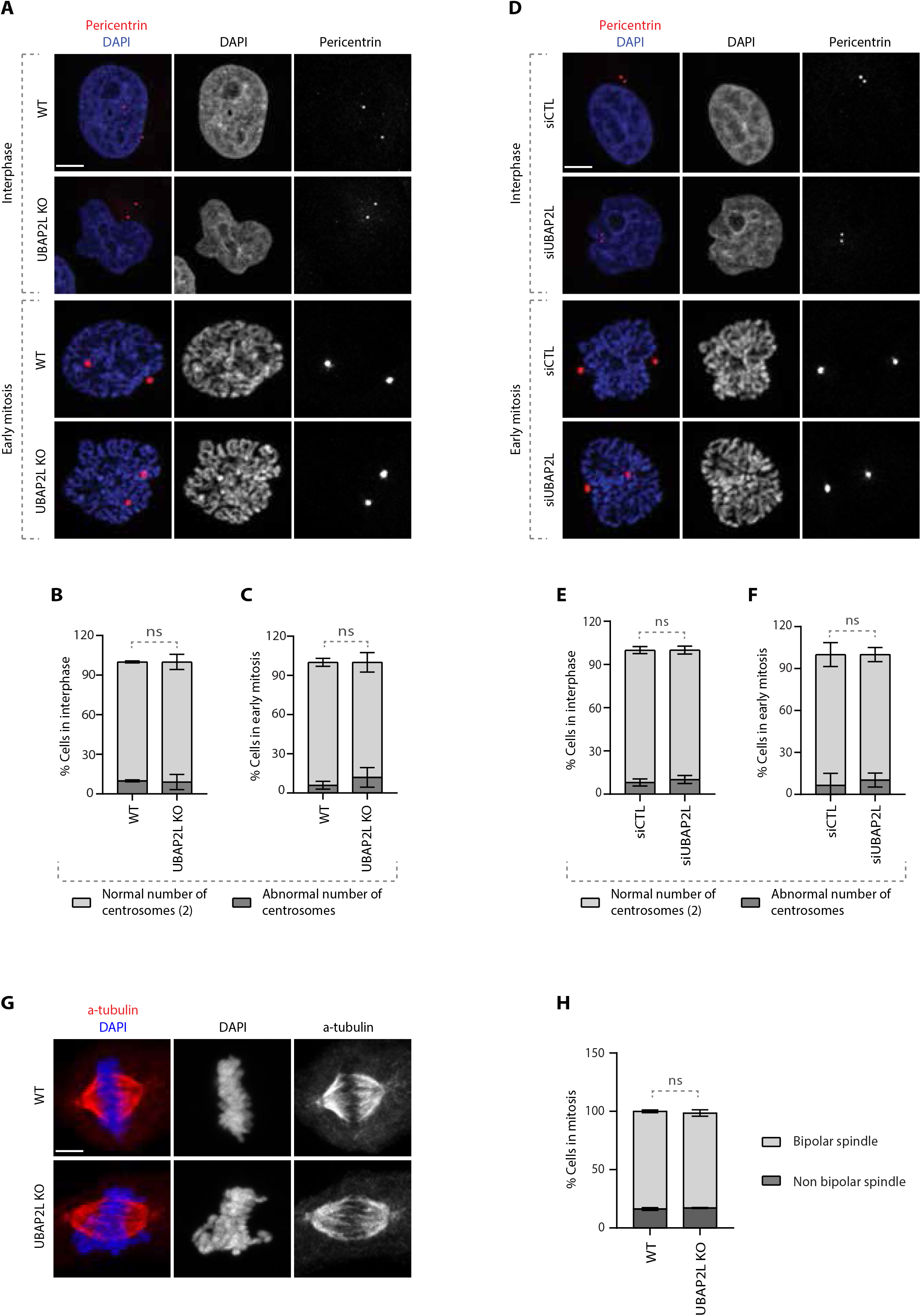
UBAP2L does not regulate spindle formation. **A-C**. Immunofluorescence (IF) representative pictures of WT or UBAP2L KO cells synchronized in G1/S with DTB or in early mitosis with DTBR **(A)** and quantification of the percentage of cells with normal (2) or abnormal number of centrosomes in interphase **(B)** or in early mitosis **(C)**. At least 50 cells per condition were analyzed for each experiment. Graphs represent the mean of three biological replicates ± SD (two sample two-tailed t-test ns=non-significant). Scale bar, 5μm. **D-F**. IF representative pictures of HeLa cells transfected with siCTL or siUBAP2L for 48h and synchronized in G1/S with DTB or in early mitosis with DTBR **(D)** and quantification of the percentage of cells with normal (2) or abnormal number of centrosomes in interphase **(E)** or in early mitosis **(F)**. At least 50 cells per condition were analyzed for each experiment. Graphs represent the mean of three biological replicates ± SD (two sample two-tailed t-test ns=non-significant). Scale bar, 5μm. **G, H**. IF representative pictures of WT or UBAP2L KO cells synchronized in mitosis with DTBR **(G)** and quantification of the percentage of cells with bipolar or non-bipolar mitotic spindle **(H)**. At least 50 cells per condition were analyzed for each experiment. The graph represents the mean of three biological replicates ± SD (two sample two-tailed t-test ns=non-significant). Scale bar, 5μm.

To better understand the role of UBAP2L in mitosis, we next characterized its subcellular localization pattern during mitotic progression. To overcome the challenge that UBAP2L is an abundant, aggregation-prone cytoplasmic protein (Youn *et al*, 2018), we applied immunofluorescence analysis to specifically extract its soluble cytoplasmic pools and we qualitatively assessed the localization of endogenous UBAP2L within different mitotic structures using appropriate protein markers (Fig EV2). Our results showed that a small fraction of UBAP2L foci may occasionally weakly co-localize to outer kinetochore subregions in prometaphase (Fig EV2A) and to inner centromere subregions in metaphase (Fig EV2B), as shown by BubR1 and INCENP labeling respectively. On the other hand, our analysis revealed that UBAP2L tends to be enriched in centrosomes and spindle poles in early mitotic stages, which was confirmed by staining with both γ-tubulin (Fig EV2C) and α-tubulin (Fig EV2D). The localization of UBAP2L at the mitotic spindle was more evident at later mitotic stages, with UBAP2L accumulating in the midzone during anaphase and in the midbody during telophase (Fig EV2E-F). Overall, our results suggest that UBAP2L is a spindle-associated protein and may exert its mitotic functions on spindle-localized downstream targets.

### UBAP2L regulates PLK1 levels and activity

Having shown that UBAP2L is required for proper mitotic progression, we next sought to address whether UBAP2L might control any essential mitotic regulators. Although UBAP2L is a predicted ubiquitin-binding protein, no downstream targets of UBAP2L have been reported so far in the context of ubiquitin signaling. Interestingly, a proteomics study in human cells suggested that UBAP2L can interact with CUL3 E3-ligase complexes (Bennett *et al*, 2010). This prompted us to investigate whether components of the mitotic CUL3 signaling (Jerabkova & Sumara, 2019) are associated with UBAP2L and its function during mitotic progression. Downregulation of UBAP2L did not affect the expression and localization of the CUL3 targets Aurora A (AurA) (Moghe *et al*, 2012) and Aurora B (AurB) (Maerki *et al*, 2009; Sumara *et al*, 2007) in mitotically arrested cells, but it increased the levels of PLK1 (Beck *et al*, 2013) (Fig EV3A). Downregulation of UBAP2L also did not affect the localization of other non-CUL3 related mitotic factors such as Cyclin B1 (Fig EV3A). Western blot analysis of cells synchronized in G1/S phase, revealed that loss of UBAP2L had no effect on the protein levels of Cyclin B1, AurA and AurB, but resulted in increased PLK1 levels compared to control cells (Fig 3A). Consistently, immunofluorescence analysis in G1/S phase showed that downregulation of UBAP2L led to an increased number of cells expressing PLK1 and that PLK1 was enriched in or near centromeres as confirmed by CREST co-staining (Fig 3B-D). These results were confirmed in two independent UBAP2L KO clones, both of which displayed increased PLK1 protein levels and enriched PLK1 peri-centromere localization during interphase (Fig 3E-H), without any obvious effects on the expression of Cyclin B1, AurA and AurB (Fig 3E and Fig EV3B-E). Subcellular fractionation assays further confirmed the nuclear accumulation of PLK1 during interphase in UBAP2L KO cells compared to WT cells (Fig 3I). The effect of UBAP2L on PLK1, prompted us to test whether UBAP2L acts on other PLK family members but no detectable changes were observed in the total protein levels of PLK2, PLK3 and PLK4 upon UBAP2L downregulation (Fig 3J). Interestingly, indirect analysis of PLK1 kinase activation as assessed by the PLK1 activatory phosphorylation at Thr210 (Macůrek *et al*, 2008) as well as by the PLK1 phospho-substrate BubR1 (Elowe *et al*, 2007) (Fig 3J and K), suggested that depletion of UBAP2L leads to increased PLK1 kinase activity in interphase. Altogether, our results suggest aberrant PLK1 signaling and increased stability of PLK1 in the absence of UBAP2L.

**Figure 3.**
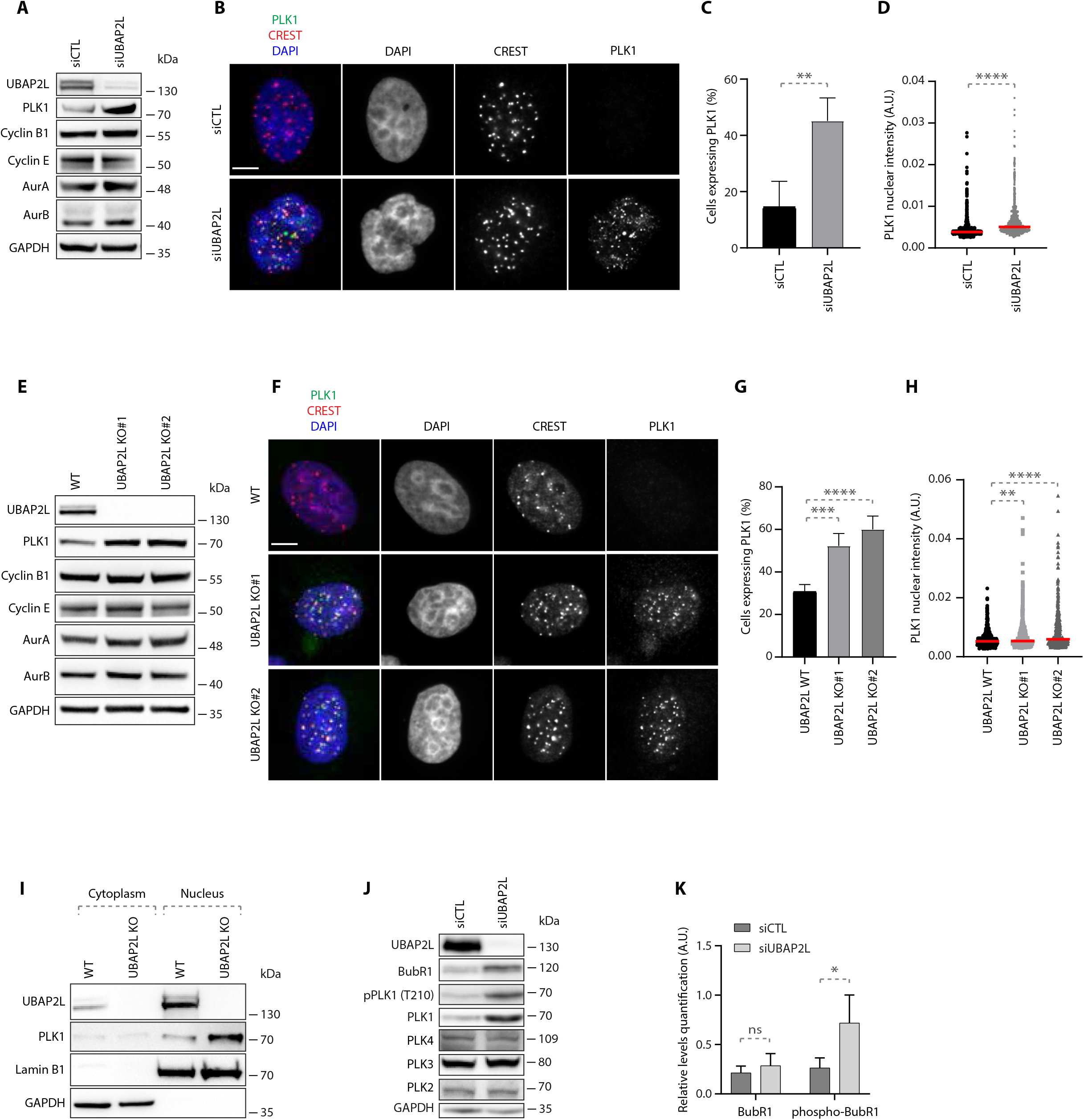
UBAP2L regulates PLK1 levels and activity. **A**. Western blot (WB) analysis of HeLa cell lysates from G1/S synchronized cells using DTB treated with siCTL or siUBAP2L. Proteins molecular weight (MW) is indicated in kilo Daltons (kDa). WB is representative of three independent replicates. **B-D**. IF representative pictures of G1/S synchronized HeLa cells treated with the indicated siRNAs and quantification of the percentage of cells expressing PLK1 **(C)** or PLK1 nuclear intensity (arbitrary units A.U.) **(D)**. Scale bar, 5μm. At least 250 cells were quantified per condition for each replicate. Graphs depicted in **(C)** represent the mean of three replicates ± SD (two sample two-tailed t-test **P<0,01). Each dot of graphs **(D)** represents PLK1 nuclear intensity in a single nucleus. The measurements of three biological replicates are combined, red bars represent the mean (Mann-Whitney test ****P<0,0001). **E**. WB analysis of WT or UBAP2L KO HeLa cell lysates from G1/S synchronized cells using DTB. Proteins MW is indicated in kDa. WB is representative of three independent replicates. **F-H**. IF representative pictures of G1/S synchronized WT or UBAP2L KO HeLa cells using DTB and quantification of the percentage of cells expressing PLK1 **(G)** or PLK1 nuclear intensity (A.U.) **(H)**. Scale bar, 5μm. At least 250 cells were quantified per condition for each replicate. Graphs depicted in **(G)** represent the mean of four replicates ± SD (one-way ANOVA with Dunnett’s correction ***P<0,001, ****P<0,0001). Each dot of graphs **(H)** represents PLK1 nuclear intensity in a single nucleus where PLK1 signal was detectable. Interphasic cells which do not express PLK1 were excluded from the quantification. The measurements of four biological replicates are combined, red bars represent the mean (Kruskal-Wallis test with Dunn’s correction **P<0,01, ****P<0,0001). **I**. WT or UBAP2L KO G1/S synchronized (DTB) HeLa cells were lysed, fractionated into cytoplasmic and nuclear fractions and analyzed by WB. Proteins MW is indicated in kDa. WB is representative of three independent replicates. **J**. WB analysis of HeLa cell lysates from unsynchronized cells treated with the indicated siRNAs. Proteins MW is indicated in kDa. WB is representative of three independent replicates. **K**. Quantification of the relative protein levels of BubR1. Graphs represent the average ratio of BubR1 (lower band) or phospho-BubR1 (upper band)/GAPDH (A.U.) from three biological replicates ± SD (two sample two-tailed t-test *P<0,05, ns=non-significant).

### UBAP2L regulates PLK1 stability in a cell-cycle specific manner

The increased PLK1 levels observed in UBAP2L KO cells could be either due to enhanced protein translation or reduced protein degradation. To distinguish between the two possibilities, we analyzed PLK1 protein levels in a time course of WT and UBAP2L KO cells treated either with the translation inhibitor cycloheximide (CHX), or with the proteasomal inhibitor MG132. In contrast to AurB, PLK1 protein levels remained stable up to 8h of CHX treatment in the absence of UBAP2L, while AurB and PLK1 were gradually degraded in WT cells, both during interphase (Fig 4A) and in cells arrested in mitosis using the microtubule stabilizing agent paclitaxel (Fig 4B). MG132 treatment increased the levels of total ubiquitin as expected but no additive effect was observed in PLK1 levels in UBAP2L depleted cells relative to WT cells (Fig 4C). Our results suggest that UBAP2L may promote proteasomal degradation of PLK1. Thus, we next aimed to determine during which cell cycle stage UBAP2L controls PLK1 stability.

**Figure 4.**
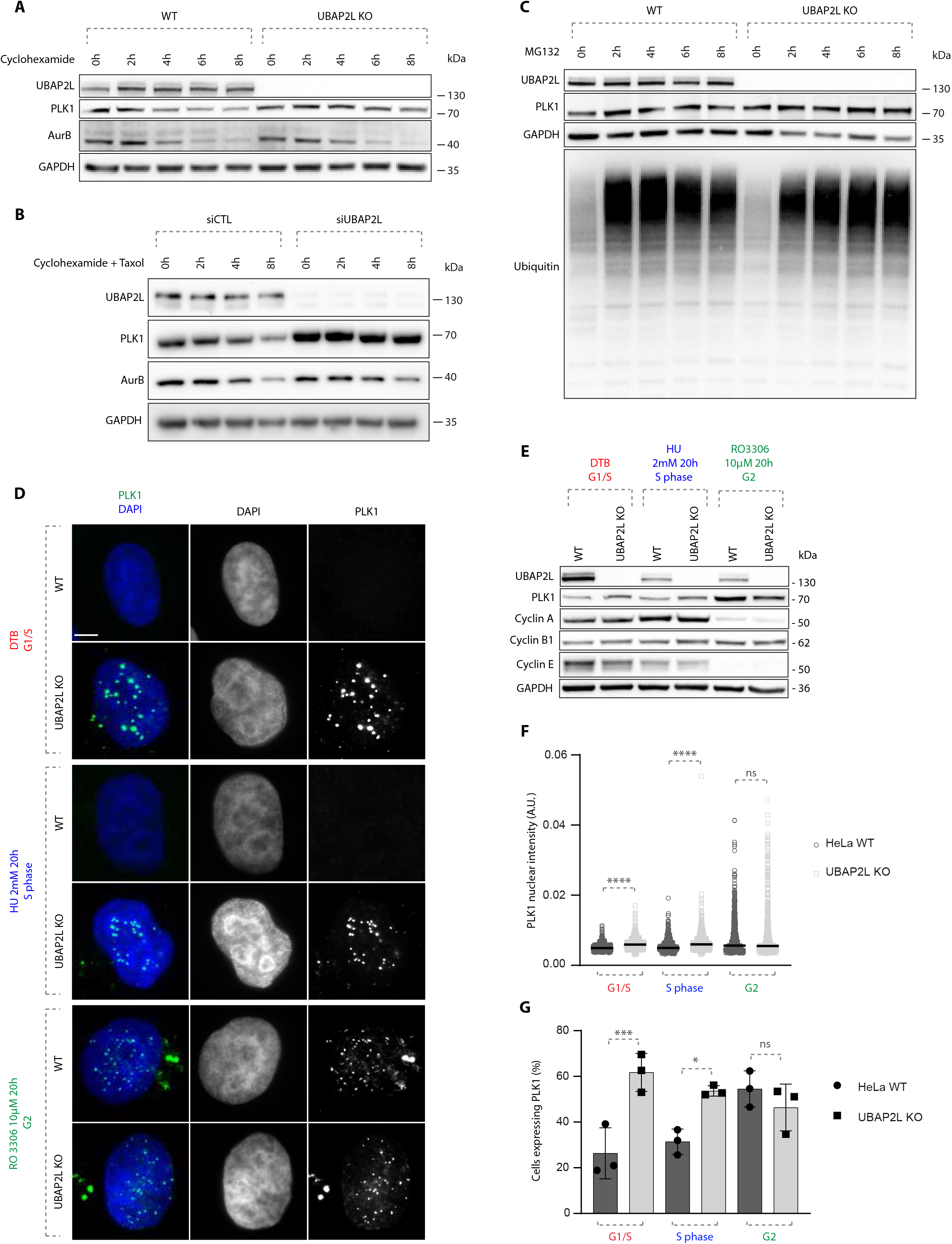
UBAP2L regulates PLK1 stability in a cell-cycle specific manner. **A**. WB analysis of WT or UBAP2L KO HeLa lysates from G1/S synchronized (DTB) cells treated with 100μg/mL cycloheximide (CHX) for the indicated times. Proteins MW is indicated in kDa. WB is representative of three independent replicates. **B**. WB analysis of control or UBAP2L-silenced HeLa lysates from mitotic cells synchronized with 1μM paclitaxel and treated with 100μg/mL CHX for the indicated times. Proteins MW is indicated in kDa. WB is representative of three independent replicates. **C**. WB analysis of WT or UBAP2L KO HeLa lysates from G1/S synchronized (DTB) cells treated with 25μM of the proteasomal inhibitor MG132 for the indicated times. Proteins MW is indicated in kDa. WB is representative of three independent replicates. **D**. Representative IF pictures of WT or UBAP2L KO HeLa cells synchronized in G1/S using DTB, in S using 2mM hydroxyurea (HU) for 20h or in G2 using 10μM of CDK1 inhibitor RO 3306 for 20h. Scale bar, 5μm. **E**. WB analysis of the experiment depicted in **(D)**. Proteins MW is indicated in kDa. WB is representative of three independent replicates. **F, G**. Quantification of PLK1 nuclear intensity (A.U.) **(F)** and of the percentage of cells expressing PLK1 **(G)**. At least 200 cells per condition were quantified for each replicate (n=3). Each dot of graphs **(F)** represents PLK1 nuclear intensity in a single nucleus. The measurements of three biological replicates are combined, red bars represent the mean. Graphs depicted in **(G)** represent the mean of three replicates ± SD. Statistics were obtained using the Mann-Whitney test **(F)** or one-way ANOVA with Dunnett’s correction **(G)** *P<0,05, ***P<0,001, ****P<0,0001, ns=non-significant.

Indeed, PLK1 protein levels strongly fluctuate during cell cycle progression, increasing in G2 phase, peaking during mitosis and decreasing again during mitotic exit and in early G1 (Golsteyn *et al*, 1995; Bruinsma *et al*, 2012). For this purpose, we analyzed PLK1 levels and localization by synchronizing cells in different cell cycle stages using several treatments: double thymidine block for G1/S transition, hydroxyurea for the S phase and CDK1 inhibitor RO3306 for G2 (Fig 4D), as previously described (Agote-Arán *et al*, 2021). These stringent synchronization protocols were used to exclude any indirect effects on PLK1 stability due to cell cycle delays in UBAP2L-deficient cells relative to WT cells. Western blot with antibodies against several cell cycle markers confirmed efficient synchronization of cells. Cyclin E accumulated during G1/S transition and decreased along the S phase, Cyclin A levels gradually increased and peaked in S phase, and Cyclin B1 gradually increased reaching the highest concentration in G2 (Fig 4E). Interestingly, the number of cells with high levels of PLK1, as well as PLK1 nuclear accumulation, were increased in UBAP2L KO cells during G1 and S phases, but no changes were detected during G2 stage compared to WT cells (Fig 4D, F and G). These results suggest that UBAP2L rather seems to regulate PLK1 levels during or after mitotic exit and not prior to mitotic entry.

### The C-terminal domain of UBAP2L mediates its function on PLK1 and on mitotic progression

Next, we aimed to understand if UBAP2L is specifically required for the regulation of PLK1 levels and localization and which domain of UBAP2L mediates its function on PLK1. Rescue experiments in UBAP2L KO cells ectopically expressing flag-tagged UBAP2L full length (FL) or UBAP2L protein fragments (Fig EV4A and B), revealed that nuclear accumulation of PLK1 in interphase could be efficiently restored by re-expression of UBAP2L FL or the UBAP2L C-terminal (CT) fragment but not the N-terminal (NT) fragment of UBAP2L (Fig 5A and B). These findings argue that the function of UBAP2L on PLK1 is mediated through its C-terminal part and it might be disconnected from the reported role of the RGG domain on mitosis (Maeda *et al*, 2016). To understand the physiological relevance of the C-terminal domain of UBAP2L during mitosis, we repeated rescue experiments in mitotically synchronized UBAP2L KO cells and assessed the presence of mitotic defects. Re-expression of UBAP2L FL or the UBAP2L CT fragment, but not the UBAP2L NT fragment, partially restored the chromosome alignment defects (Fig 5C and D) and the chromosome segregation errors (Fig 5D and EV3C) in UBAP2L depleted cells. Moreover, since loss of UBAP2L frequently led to mitotic cell death (Fig 1A, F, H, M and Movies EV3 and EV6), we tested whether UBAP2L might also regulate cell proliferation. In accordance with studies showing that cells harboring accumulated errors during cell division are often characterized by reduced survival (Cheng & Crasta, 2017), UBAP2L KO cells displayed significantly reduced long-term proliferation capacity and viability (Fig 5E-G). Re-expression of UBAP2L FL or the UBAP2L CT fragment but not the UBAP2L NT fragment partially rescued cell proliferation (Fig 5E and F) and fully rescued cell survival (Fig 5E and G). In view of the partial rescue of mitotic defects and cell proliferation by UBAP2L FL and UBAP2L CT, we cannot exclude that the UBAP2L-dependent regulation of mitosis may not be restricted to the control of PLK1 stability, but it may also involve additional, yet unidentified mitotic factors.

**Figure 5.**
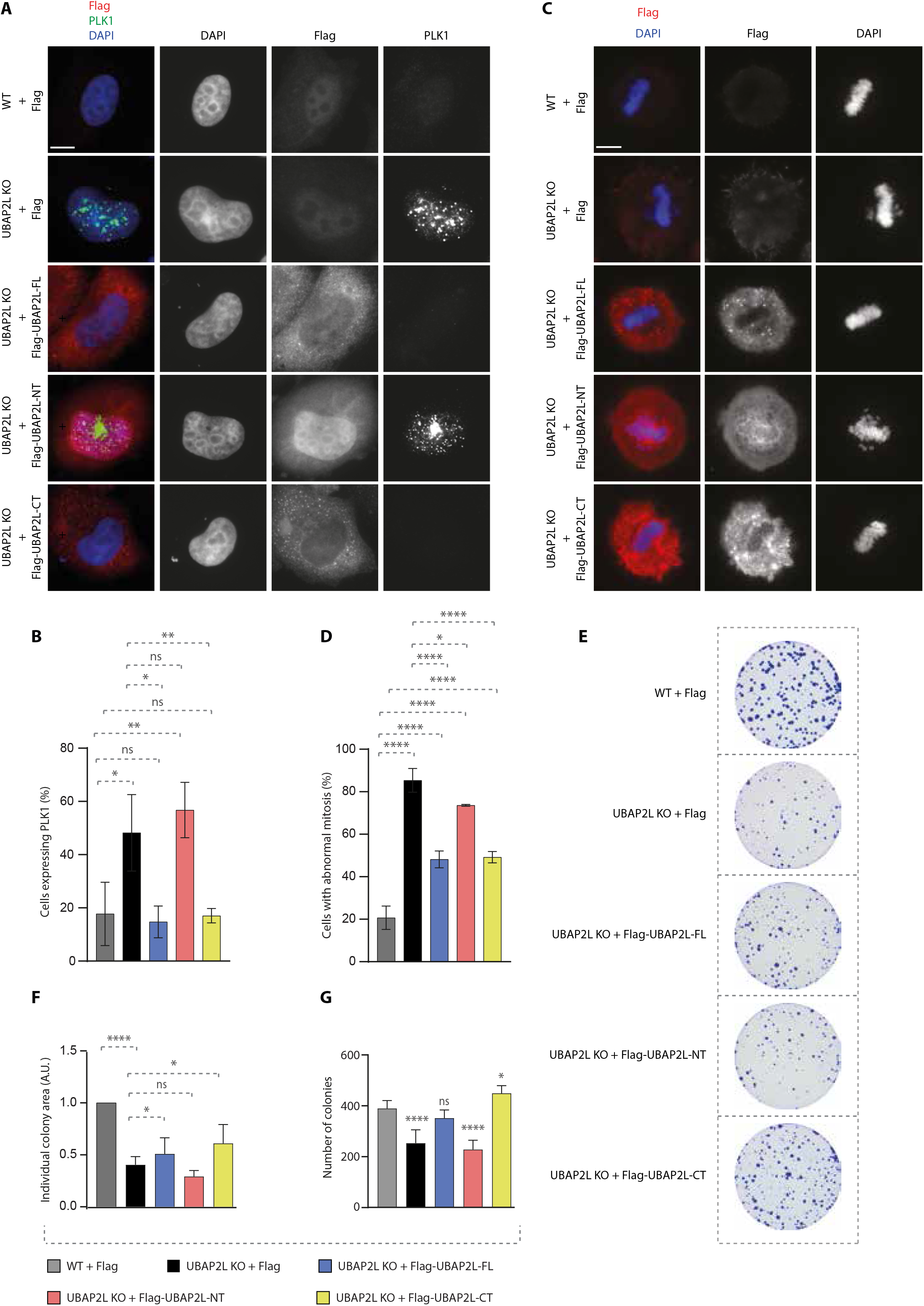
The C-terminal domain of UBAP2L mediates its function on PLK1 and on mitosis. **A, B**. IF analysis of G1/S synchronized WT or UBAP2L KO HeLa cells using DTB and transfected with the indicated flag-tagged UBAP2L protein fragments for 48h **(A)** and quantification of the percentage of cells expressing PLK1 **(B)**. Scale bar, 5μm. At least 100 cells per condition were quantified for each experiment. Graphs represent the mean of three replicates ± SD (one-way ANOVA with Sidak’s correction *P<0,05, **P<0,01, ns=non-significant). **C, D**. IF representative pictures of WT or UBAP2L KO HeLa cells transfected with the indicated flag-tagged UBAP2L protein fragments for 48h and synchronized in mitosis using monastrol release (MR) **(C)** and quantification of the percentage of cells with abnormal mitosis (misalignments and/or DNA bridges) **(D)**. Scale bar, 5μm. At least 50 cells per condition were quantified for each experiment. Graphs represent the mean of three replicates ± SD (one-way ANOVA with Sidak’s correction *P<0,05, ****P<0,0001). **E-G**. Colony formation assay of WT or UBAP2L KO HeLa cells transiently transfected with the indicated flag-tagged UBAP2L protein fragments and quantification of the individual colony area **(F)** and of the number of colonies **(G)** after 7 days of culture. Graphs represent the mean of three replicates ± SD (one-way ANOVA with Sidak’s correction *P<0,05, ****P<0,0001, ns=non-significant).

### UBAP2L-mediated regulation of PLK1 is independent of stress signaling

The Domain of Unknown Function (DUF) located within the C-terminal part of UBAP2L is responsible for its interaction with core components of stress granules (SGs) such as the Ras GTPase-activating protein-binding protein (G3BPs), thus enabling their correct assembly upon stress signaling (Huang *et al*, 2020). The fact that UBAP2L exerts its function on PLK1 via the C-terminal, rather than via its N-terminal part which is associated with the ubiquitin-binding properties of UBAP2L, prompted us to investigate whether UBAP2L-mediated regulation of PLK1 is linked to SG signaling. To this end, we downregulated G3BP1 and G3BP2 by specific siRNAs (Cirillo *et al*, 2020) (Fig EV4D) in interphasic UBAP2L KO cells and tested whether UBAP2L re-expression could still restore PLK1 accumulation. G3BPs depletion alone in UBAP2L KO cells, led to moderately reduced number of cells expressing PLK1 in interphase (Fig 6A-B), as well as to decreased PLK1 total protein levels and nuclear intensity (Fig EV4D and E). However, down-regulation of G3BPs did not interfere with the ability of UBAP2L FL and CT part to reverse the elevated PLK1 levels and nuclear accumulation in interphase observed in the UBAP2L KO cells (Fig 6A-B and EV4E). Thus, our results suggest that UBAP2L-mediated regulation of PLK1 can be uncoupled from the previously established function of the UBAP2L C-terminal domain in G3BP1/G3BP2-dependent SGs assembly.

**Figure 6.**
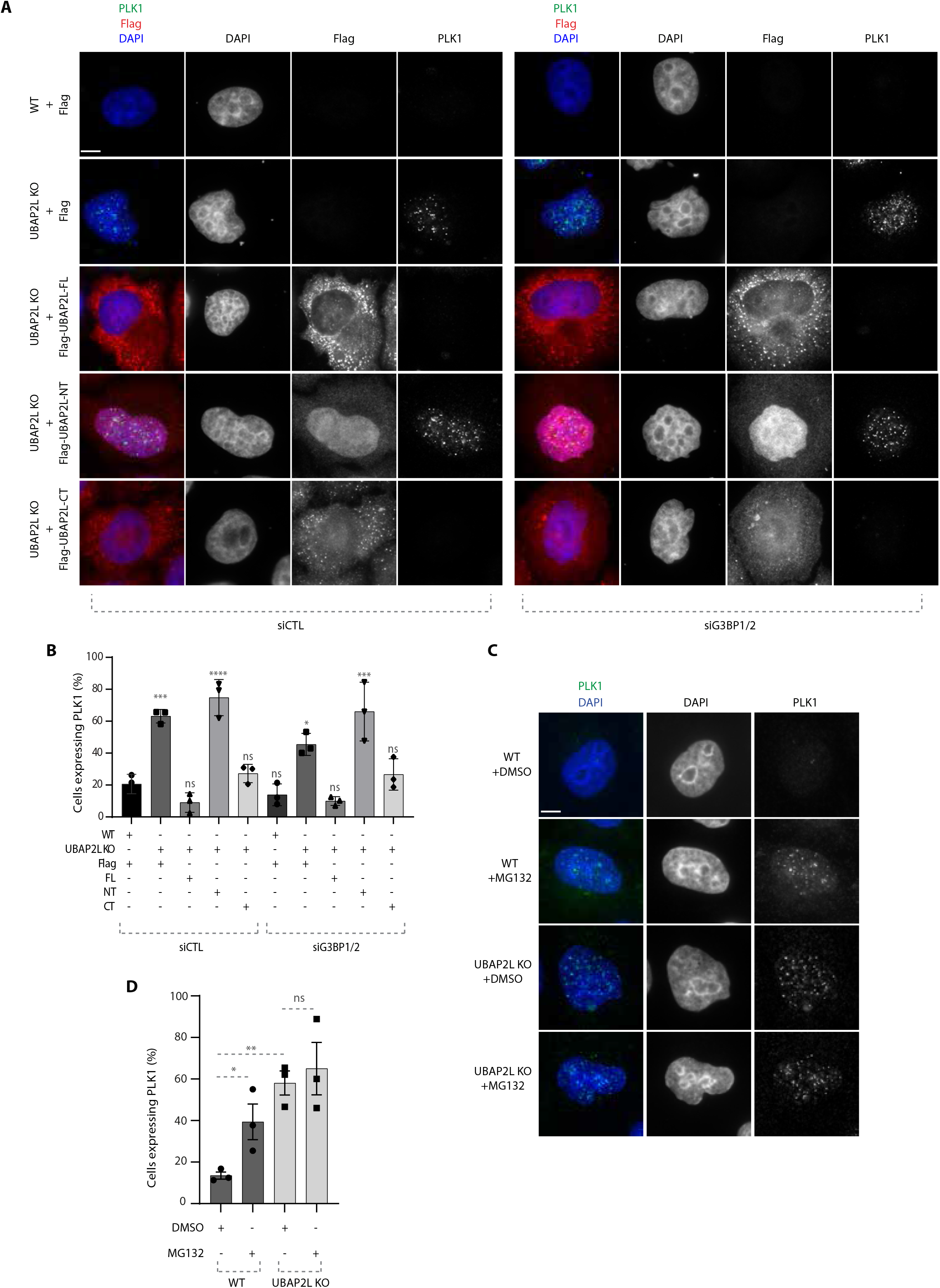
UBAP2L-mediated regulation of PLK1 is independent of stress signaling. **A, B**. Representative IF images of WT or UBAP2L KO HeLa cells synchronized in G1/S using DTB and transfected with the indicated flag-tagged UBAP2L constructs and control or G3BP1/2 siRNAs for 48h **(A)** and quantification of the percentage of cells expressing PLK1 **(B)**. Scale bar, 4μm. At least 150 cells were quantified per condition for each experiment. The graph depicted in **(B)** represents the mean of three replicates ± SD (one-way ANOVA with Sidak’s correction *P<0,05, ***P<0,001, ****P<0,0001, ns=non-significant). **C, D**. Representative IF images of WT or UBAP2L KO HeLa cells synchronized in G1/S using DTB and treated with vehicle (DMSO) or 25μM of MG132 for 4h **(C)** and quantification of the percentage of cells expressing PLK1 **(D)**. Scale bar, 4μm. At least 150 cells were quantified per condition for each experiment. The graph depicted in **(D)** represents the mean of three replicates ± SD (one-way ANOVA with Sidak’s correction *P<0,05, **P<0,01, ns=non-significant).

Having excluded the involvement of SG signaling in the UBAP2L-mediated PLK1 regulation during G1/S phase, we wondered whether other types of stress might be implicated in our pathway. Therefore, we induced proteotoxic stress by MG132 treatment and compared the levels and localization of PLK1 in WT versus UBAP2L KO cells. MG132 treatment alone led to increased number of G1/S synchronized cells with elevated PLK1 levels in the nucleus (Fig 6C-D). However, the effect of MG132 on PLK1 stability was milder compared to depletion of UBAP2L and most importantly it was not additive (Fig 6C-D). We thus conclude that UBAP2L-mediated regulation of PLK1 may occur independently of stress signaling.

### UBAP2L controls PLK1 turnover during mitotic progression

Our results so far suggest that the effect of UBAP2L on PLK1 stability is cell-cycle specific and seems to occur after mitotic entry (Fig 4). This observation triggered us to investigate in more detail how UBAP2L regulates the spatiotemporal dynamics of PLK1 throughout mitotic progression. To this end, we assessed the effect of UBAP2L depletion on the stability of PLK1 in mitotically synchronized cells treated with monastrol and collected at different time points after being released from the treatment. Immunofluorescence analysis revealed that as early as in prometaphase, PLK1 displayed increased levels upon UBAP2L depletion compared to control cells (Fig 7A). Moreover, during telophase and cytokinesis stages (1h and 30 min post release), UBAP2L depletion not only led to the enrichment of PLK1 in the midbody, but PLK1 was also aberrantly retained at the kinetochores compared to control cells (Fig 7A-C). Finally, when UBAP2L depleted cells exited mitosis and entered into the subsequent interphase (3h, 4 h and 30min post release), PLK1 was still highly enriched in the regions of kinetochores compared to control cells in which PLK1 was no longer detectable at this subcellular location (Fig 7A and B). Interestingly and consistent with our previous results on PLK1 stability in G1 and in prometaphase arrested cells (Fig 4A and B), UBAP2L downregulation led to increased PLK1 stability after release from monastrol (Fig 7D). To further corroborate these findings, we performed live video imaging in HeLa PLK1-eGFP knock in (KI) cells (Fig EV5A-E) that were synchronized with double thymidine block and release in the presence or absence of UBAP2L. Indeed, loss of UBAP2L led to aberrant accumulation of PLK1 from early prophase to late cytokinesis in several mitotic structures including kinetochores, spindle poles, midzone and midbody (Fig 7E and Movies EV7 and EV8). Altogether, our results suggest that UBAP2L is required for the efficient degradation of PLK1 and its removal from different mitotic structures during mitotic exit.

**Figure 7.**
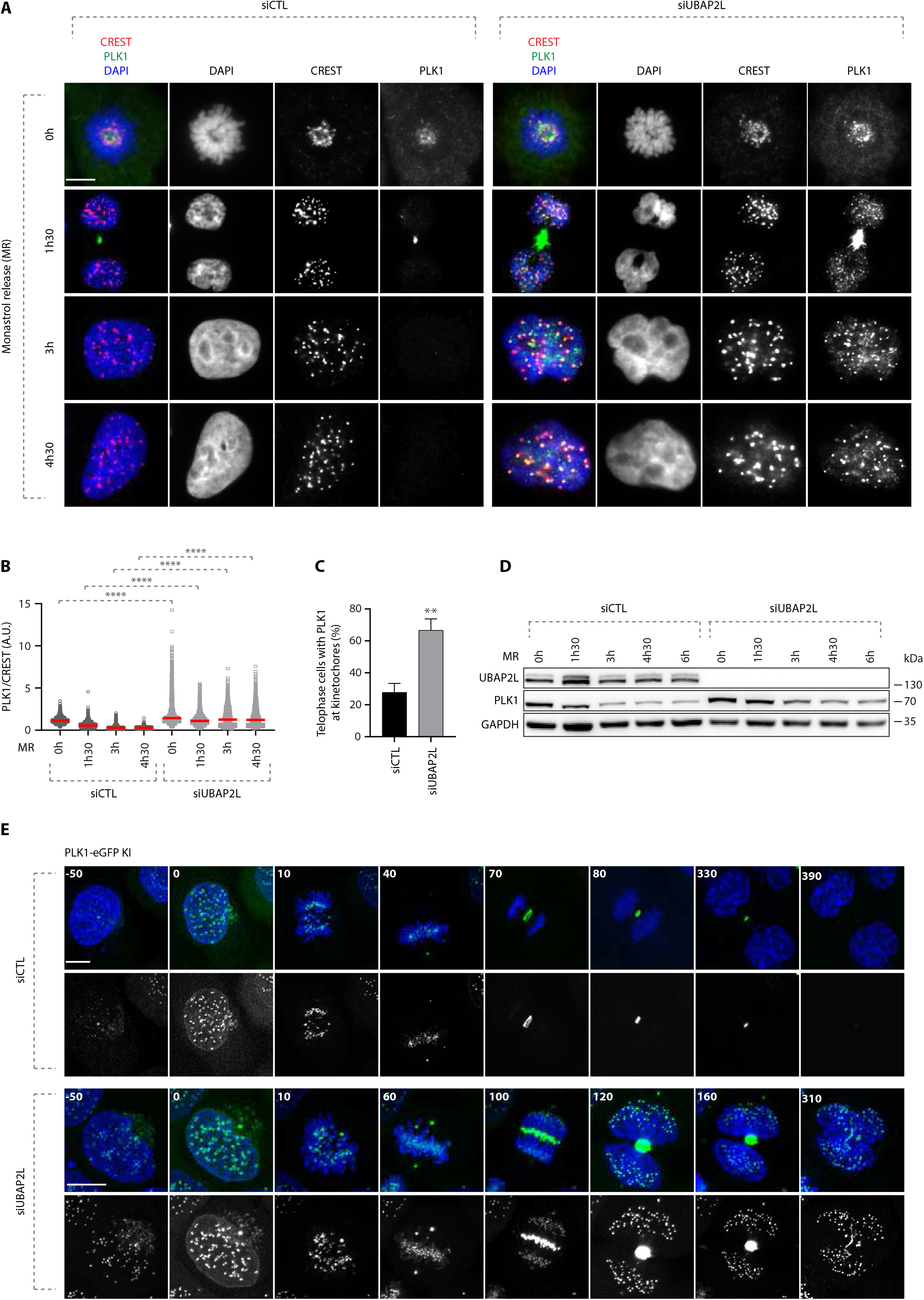
UBAP2L controls PLK1 turnover during mitotic progression. **A-C**. Representative IF images of control or UBAP2L-downregulated cells synchronized in prometaphase using 1mM monastrol and released at the indicated time points. Scale bar, 5μm. Quantification of PLK1 relative kinetochores intensity (A.U.) **(B)** and of the percentage of telophase cells with PLK1 at kinetochores **(C)**. At least 250 cells per condition were quantified for each replicate. Each dot of **(B)** represents PLK1/CREST intensity ratio at a single pair of kinetochores. At least 10 pairs of kinetochores per cell were analyzed and the measurements of three biological replicates are combined, red bars represent the mean (Kruskal-Wallis test with Dunn’s correction ****P<0,0001). Graph **(C)** represents the mean of three replicates ± SD (two sample two-tailed t-test **P<0,01). **D**. WB analysis of control or UBAP2L-downregulated cell lysates after MR at the indicated time points. Proteins MW is indicated in kDa. WB is representative of three independent replicates. **E**. Spinning disk time-lapse microscopy of PLK1-eGFP Knock-In (KI) HeLa cells synchronized with DTBR in mitosis. The selected frames of the movies are depicted and the corresponding time is indicated in minutes. SiR-DNA was used for DNA staining. Scale bars, 5μm and 8μm.

To better characterize the effects of UBAP2L depletion on PLK1 turnover, we analyzed in more detail the individual mitotic pools of PLK1. To this end, we synchronized WT and UBAP2L KO cells with double thymidine block and release and compared the localization of endogenous PLK1 within different mitotic structures using appropriate protein markers (Fig 8). As early as in prometaphase, the chromosome-bound fraction of PLK1 increased, with PLK1 being enriched in both outer kinetochore (Fig EV5F and G) and inner centromere (Fig 8A and B) subregions, as revealed by analysis of BubR1 and INCENP localization, respectively. PLK1 was also enriched in centrosomes during prometaphase and metaphase as confirmed by pericentrin staining (Fig 8C and D). Moreover, in UBAP2L depleted cells PLK1 was significantly enriched in the midbody during telophase/cytokinesis, as shown by PRC1 labelling (Fig 8E and F). Finally, UBAP2L depletion led to the accumulation of the spindle-bound fraction of PLK1 around centrosomes and at the midzone region as revealed by α-tubulin staining (Fig 8G). Overall, we conclude that UBAP2L acts as a major regulator of PLK1 stability and controls globally the protein turnover of PLK1 within distinct mitotic structures.

**Figure 8.**
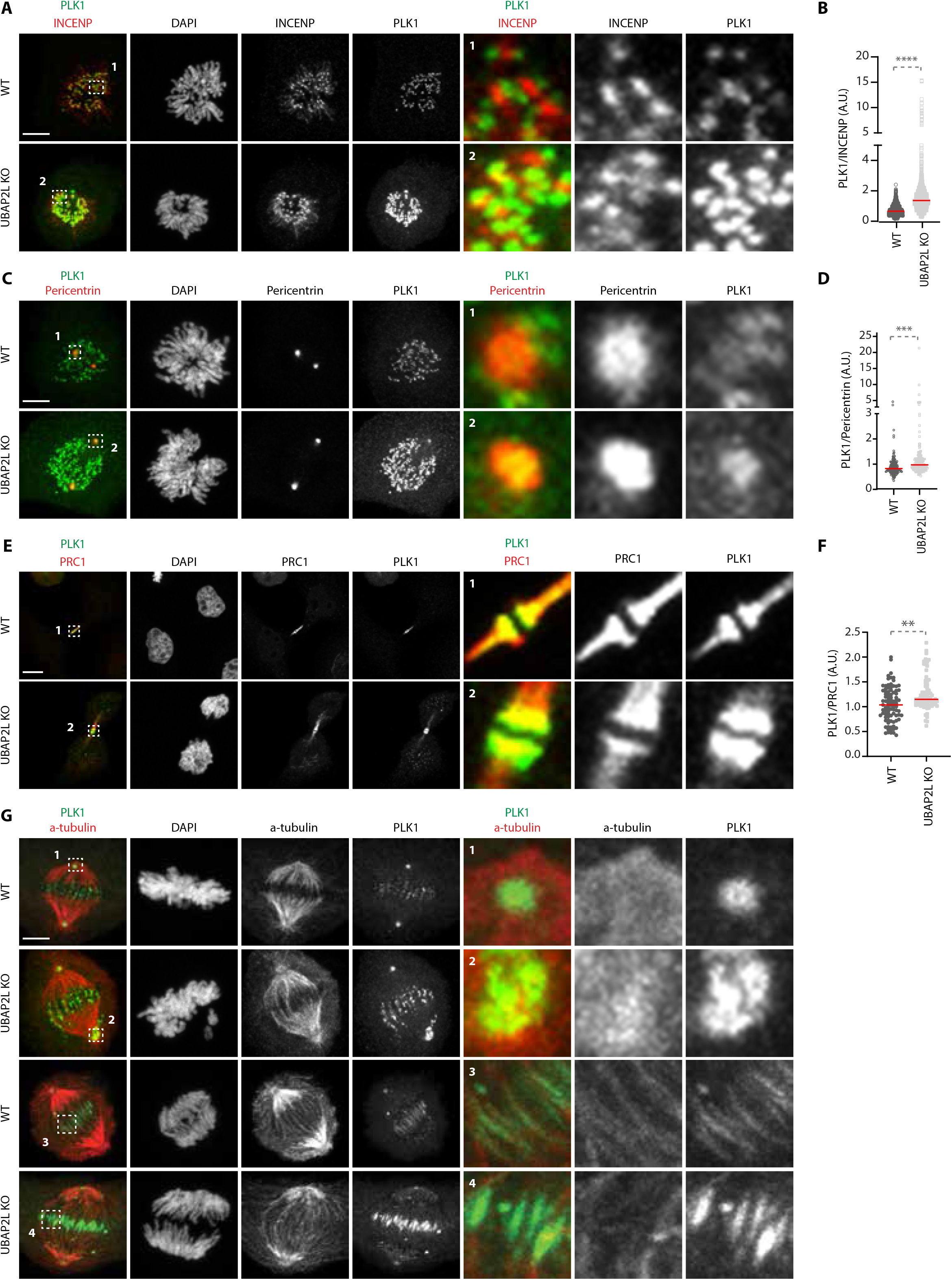
UBAP2L controls PLK1 localization at several mitotic structures. **A, B**. Representative IF pictures of WT or UBAP2L KO HeLa cells synchronized in mitosis with DTBR **(A)** and quantification of the relative PLK1 intensity at the inner centromere (A.U.) **(B)**. ROIs are shown in the corresponding numbered panels. Scale bar, 5μm. At least 50 cells were quantified per condition for each experiment. Each dot represents PLK1/INCENP intensity ratio at a single centromere. The measurements of three biological replicates are combined, red bars represent the mean (Mann-Whitney test ****P<0,0001). **C, D**. Representative IF pictures of WT or UBAP2L KO HeLa cells synchronized in mitosis with DTBR **(C)** and quantification of the relative PLK1 intensity at the centrosomes (A.U.) **(D)**. ROIs are shown in the corresponding numbered panels. Scale bar, 5μm. At least 50 cells were quantified per condition for each experiment. Each dot represents PLK1/Pericentrin intensity ratio at a single centrosome. The measurements of three biological replicates are combined, red bars represent the mean (Mann-Whitney test ****P<0,0001). **E, F**. Representative IF pictures of WT or UBAP2L KO HeLa cells synchronized in mitosis with DTBR **(E)** and quantification of the relative PLK1 intensity at the midbody (A.U.) **(F)**. ROIs are shown in the corresponding numbered panels. Scale bar, 4μm. At least 50 cells were quantified per condition for each experiment. Each dot represents PLK1/PRC1 intensity ratio at the midbody. The measurements of three biological replicates are combined, red bars represent the mean (Mann-Whitney test **P<0,01). **G**. Representative IF pictures of WT or UBAP2L KO HeLa cells synchronized in mitosis with DTBR (n=3) to qualitatively assess the localization of PLK1 at the mitotic spindle with α-tubulin staining. ROIs are shown in the corresponding numbered panels. Scale bar, 5μm.

### Increased stability of PLK1 may cause mitotic defects in UBAP2L depleted cells

Our results so far support that UBAP2L controls PLK1 levels possibly through ubiquitin-mediated degradation. However, the ubiquitin machinery components involved in the UBAP2L-dependent signaling remain unknown. Considering that UBAP2L is reported to interact with CUL3 complexes in human cells (Bennett *et al*, 2010) and that PLK1 is mono-ubiquitylated by CUL3 (Beck *et al*, 2013), we tested the involvement of UBAP2L in the CUL3/PLK1 pathway. Co-immunoprecipitation (co-IP) assays in mitotically synchronized cells showed that endogenous UBAP2L could efficiently interact with CUL3 and its substrate specific adaptor KLHL22 as well as with PLK1, relative to IgG control, but not with AurB which is another known mitotic ubiquitylation substrate of CUL3 (Beck et al., 2013; Courtheoux et al., 2016; Krupina et al., 2016; Maerki et al., 2009; Sumara et al., 2007) (Fig 9A). Furthermore, endogenous co-IP of PLK1 in mitotically synchronized cells showed that UBAP2L depletion reduced the interaction of PLK1 with CUL3 relative to control cells expressing UBAP2L (Fig 9B), indicating that UBAP2L may be an essential component of this pathway. However, since CUL3-mediated mono-ubiquitylation of PLK1 does not affect the protein stability of PLK1 (Beck *et al*, 2013), we next aimed at understanding if the polyubiquitylation status of PLK1 can be also regulated by UBAP2L. To this end, we performed co-IP of GFP-PLK1 in the presence of proteasomal inhibitor MG132 under denaturing conditions in UBAP2L KO and WT cells. Interestingly, we observed reduced polyubiquitin modification on immunoprecipitated GFP-PLK1 in cells depleted for UBAP2L (Fig 9C), suggesting that UBAP2L is additionally involved in the ubiquitin-mediated proteolysis of PLK1 during mitotic exit. The identity and precise mechanism of the possible additional E3-ligase/s involved in UBAP2L regulation of PLK1 stability remains to be determined in the future.

**Figure 9.**
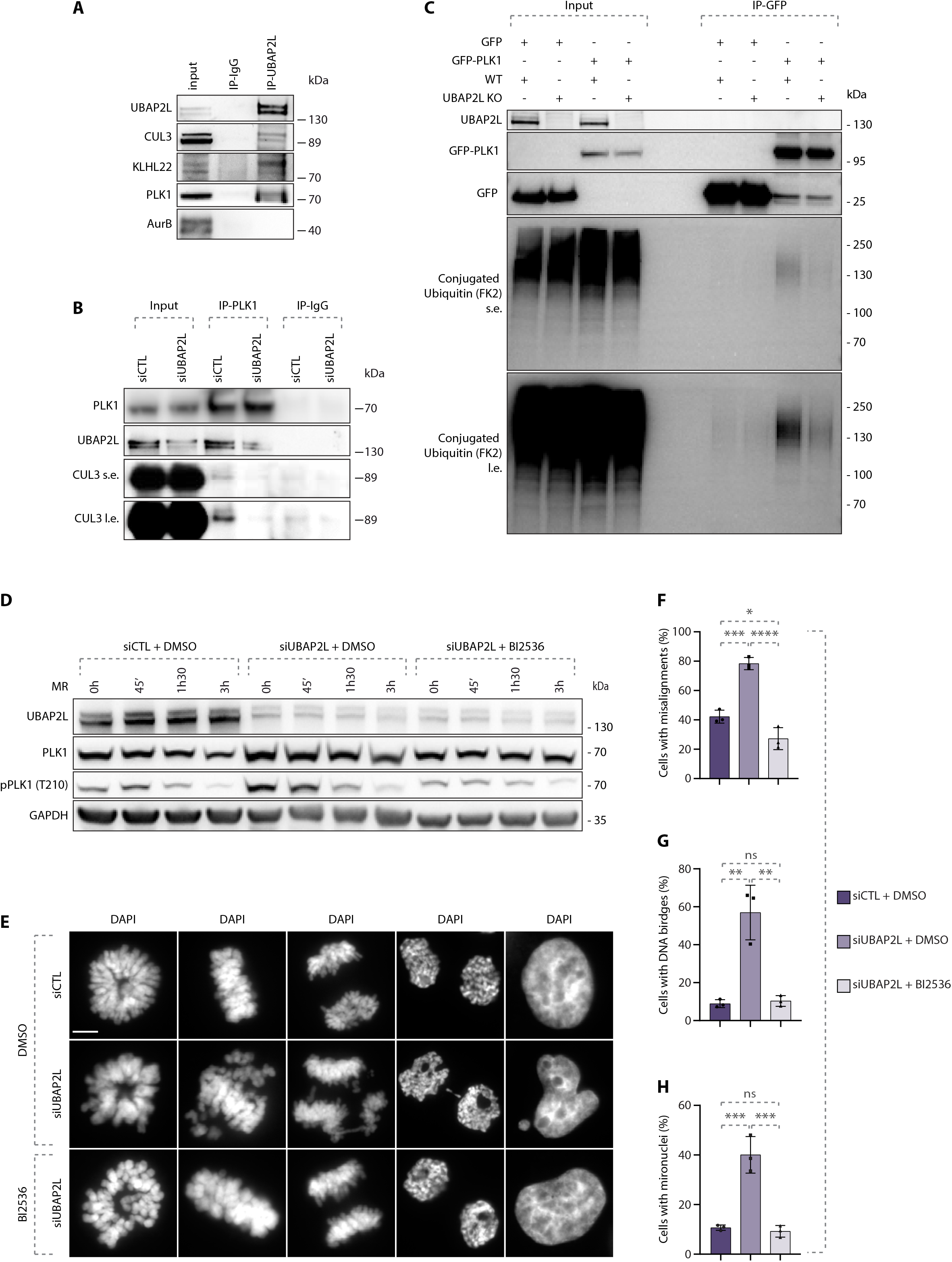
Increased stability of PLK1 may cause mitotic defects in UBAP2L depleted cells. **A**. WB analysis of endogenous immunoprecipitation (IP) of IgG or UBAP2L from HeLa cells synchronized in mitosis using 5μM STLC for 16h. Proteins MW is indicated in kDa. WB is representative of three independent replicates. **B**. WB analysis of endogenous IP of IgG or PLK1 from HeLa cells transfected with control or UBAP2L siRNA for 48h and synchronized in mitosis using 5μM STLC for 16h. Proteins MW is indicated in kDa. WB is representative of three independent replicates. **C**. WB analysis of IP under denaturing conditions of WT or UBAP2L KO HeLa cells transfected with plasmids encoding for GFP-PLK1 and His-Ubiquitin for 30h and synchronized in mitosis using 1μM Paclitaxel for 16h. The short exposure (s.e.) and long exposure (l.e.) of the membrane blotted against the FK2 antibody that specifically recognizes conjugated but not free ubiquitin are shown. Proteins MW is indicated in kDa. WB is representative of three independent replicates. **D**. WB analysis of control (siCTL) or siUBAP2L 48h-treated HeLa cells were synchronized with 1mM monastrol for 16h, treated with DMSO or with 10nM of the PLK1 inhibitor BI2536 for 45min, subsequently washed out from monastrol for the indicated time and collected for protein extraction. This moderate treatment is sufficient to restore the aberrant PLK1 catalytic activity observed in UBAP2L KO cells to the levels of the WT, enabling correct mitotic progression but prevents control cells to progress through mitosis. Proteins MW is indicated in kDa. WB is representative of three independent replicates. **E-H**. DAPI staining of the experiment described in **(D)** showing different mitotic stages **(E)**. Quantification of the percentage of cells with misalignments **(F)**, DNA bridges **(G)** and micronuclei **(H)**. Scale bar, 5μm. At least 100 cells from each mitotic stage were quantified for all conditions. Graphs represent the mean of three replicates ± SD (one-way ANOVA with Sidak’s correction *P<0,05, **P<0,01, ***P<0,001, ****P<0,0001, ns=non-significant).

Nevertheless, we asked the question whether the mitotic defects observed in cells lacking UBAP2L could be directly linked to increased PLK1 stability and aberrant PLK1 signaling. We thus performed rescue experiments in which we inhibited PLK1 activity after release from monastrol treatment using small doses of the chemical inhibitor BI2536 (Lenart *et al*, 2007) and we compared the rate of segregation errors in UBAP2L-downregulated cells relative to control cells. Indirect analysis of PLK1 kinase activation as assessed by the PLK1 activatory phosphorylation at Thr210 (Macůrek *et al*, 2008), verified that BI2536 efficiently restored PLK1 activity to control levels in UBAP2L depleted cells (Fig 9D). Interestingly, BI2536 treatment fully rescued all types of erroneous mitotic phenotypes observed in UBAP2L depleted cells, including chromosome misalignment in metaphase, DNA bridges in anaphase/telophase and micronuclei formation after cytokinesis completion (Fig 9E-H). Our results suggest that aberrant PLK1 activity resulting from increased stability of this kinase may represent the leading cause for mitotic defects observed in UBAP2L depleted cells.

## Discussion

In summary, our study describes a novel UBAP2L-mediated mechanism that spatiotemporally controls PLK1 turnover during mitotic progression. We demonstrate that UBAP2L depleted cells are characterized by significant mitotic delay, severe segregation errors and micronuclei formation but loss of UBAP2L has no effect on mitotic spindle polarity and centrosome number (Fig 2). We propose that these phenotypes can be linked to aberrant PLK1 signaling (Fig 1 and 9), as in the absence of UBAP2L PLK1 is abnormally retained at several mitotic structures and fails to get degraded during mitotic exit (Fig 7 and 8). As a result, the elevated PLK1 levels and kinase activity ultimately cause genomic instability and cell death. Importantly, we show that increased stability of PLK1 may cause mitotic defects in UBAP2L depleted cells, which could be rescued upon chemical inhibition of PLK1 (Fig 9). Thus, our data suggest that UBAP2L is required to finetune the ubiquitin-mediated PLK1 turnover during mitosis as a means to maintain genome fidelity.

### How does UBAP2L regulate mitosis?

Mitosis is a fundamental process in eukaryotes, where steps such as chromosome congression and alignment need to be precisely finetuned to ensure high fidelity of cell division (McIntosh, 2016). Phosphorylation and ubiquitylation pathways are tightly interconnected during mitosis, however how exactly these signaling cues are integrated and orchestrated in a space and time-dependent manner is still not fully understood. Here, we identify the ubiquitin-binding protein UBAP2L as an important regulator of PLK1 kinase and of faithful chromosome segregation. We show that UBAP2L regulates PLK1 in a cell-cycle specific manner, with UBAP2L depletion leading to increased PLK1 levels in mitosis and in the subsequent G1/S, while we do not observe any effect during G2 (Fig 4). UBAP2L may exert a dual role on PLK1 throughout mitotic progression, by coordinating both its proper dynamics at centrosomes, kinetochores and mitotic spindle (Fig 7 and 8), as well as its protein stability (Fig 4). UBAP2L regulates PLK1 but does not affect other PLK family members (Fig 3J), nor other components of the mitotic CUL3 signaling network such as AurA and AurB (Fig3 and EV3) that we tested so far, but it cannot be formally excluded that other targets of UBAP2L exist during mitosis. Interestingly, a pool of UBAP2L localizes to spindle-associated structures such as the centrosomes, midzone and midbody (Fig EV2). We could therefore speculate that the mitotic role of UBAP2L can be attributed to its microtubule/spindle-associated fraction which provides contacts to several subcellular structures where mitotic factors, such as PLK1, tend to localize to. Therefore, it would be worth investigating whether additional spindle and centrosome proteins might be under UBAP2L regulation to ensure mitotic fidelity. Of interest, UBAP2L has also been proposed to be phosphorylated during mitosis (Dephoure *et al*, 2008; Maeda *et al*, 2016), but the kinase involved or the underlying mechanisms are currently unknown. Given that the C-terminal domain of UBAP2L is predicted to harbor several PLK1 consensus motifs (Santamaria *et al*, 2011), it would be interesting to address the possibility of UBAP2L being a direct phosphorylation target of PLK1 or an indirect substrate via CDK1 priming phosphorylation (Parrilla *et al*, 2016). Such a regulatory feedback loop has already been described for PLK1/USP16 (Zhuo *et al*, 2015, 16) and would advance our understanding on how PLK1 can dynamically drive its own turnover to ensure fidelity of cell division.

### Role of UBAP2L C-terminal domain in the regulation of PLK1

Our data demonstrate that the uncontrolled enrichment of PLK1 in mitotic structures and its elevated protein stability during mitotic exit in UBAP2L depleted cells are specifically mediated via UBAP2L. The observed PLK1 phenotypes are corroborated by specific siRNAs against UBAP2L (Cirillo *et al*, 2020) and by CRISPR-mediated genetic depletion of UBAP2L, excluding the possibility of an off-target or a compensatory effect. Our rescue experiments provide evidence that both PLK1 aberrant accumulation and cell survival can be entirely rescued by overexpression of the C-terminal domain of UBAP2L but are not dependent on its N-terminal domain (Fig 5), that was until now considered to mediate the mitotic role of UBAP2L (Maeda *et al*, 2016). However and in line with our results, the study by Maeda and colleagues reported that an extra sequence after the UBA-RGG domain is essential for proper mitotic progression, while overexpression of the UBA-RGG domain alone cannot restore the multinuclear phenotype observed in UBAP2L depleted cells (Maeda *et al*, 2016). Nevertheless, given that mitotic defects and cell proliferation in UBAP2L depleted cells are only partially rescued by overexpression of the C-terminal domain of UBAP2L (Fig 5 and EV4), we cannot exclude the existence of additional mitotic pathways downstream of UBAP2L that do not rely on PLK1. This partial rescue of mitotic defects could also be explained by the fact that UBAP2L forms aggregates when overexpressed, suggesting that it might not localize properly or that it can lead to sequestration of essential mitotic factors.

UBAP2L and in particular its C-terminal domain have been mostly studied in the context of SGs signaling (Cirillo *et al*, 2020; Huang *et al*, 2020; Youn *et al*, 2018). Our results show that depletion of core SGs components had no effect on the ability of UBAP2L FL and/or UBAP2L CT fragment to fully restore the aberrant accumulation of PLK1 (Fig 6 and EV4). This suggests that the C-terminus domain of UBAP2L has an unexpected new role during mitosis that seems unrelated to its established role in G3BP1/G3BP2-dependent SGs signaling. Furthermore, the PRMT1-dependent UBAP2L methylation that is linked to accurate chromosome segregation, was recently reported to impair SG assembly (Huang *et al*, 2020), again indicating that the role of UBAP2L in mitosis and its effect on PLK1 does not interfere with its role is SGs signaling. Interestingly, SGs cannot be formed during mitosis and most of the known membraneless organelles are dissolved at G2/M transition in a kinase-dependent manner (Rai *et al*, 2018). It would be intriguing to speculate that UBAP2L may be subjected to phosphorylation during mitotic entry as a means to promote the dissolution of SGs, thereby shifting the interactions and functions of UBAP2L towards components of the mitotic machinery.

### How exactly does UBAP2L regulate PLK1 to ensure fidelity of cell division?

Our data demonstrate that in cells lacking UBAP2L, PLK1 not only persists at several mitotic structures from early prometaphase until late cytokinesis (Fig 8 and EV5) but is also protected from degradation (Fig 4 and 7), resulting in persistent PLK1 protein stability and activity in interphasic cells. More specifically, we show that in the absence of UBAP2L, PLK1 is resistant to CHX treatment both in interphasic (Fig 4A) and mitotic cells (Fig 4B) and that the number of cells with high PLK1 levels during interphase is significantly increased compared to control WT cells where PLK1 is only detected at basal levels (Fig 3). Furthermore, we show that in UBAP2L depleted cells PLK1 accumulates in kinetochores, centromeres, centrosomes, midzone and midbody (Fig 8 and EV5). Finally, we observe that loss of UBAP2L weakens the mitotic interaction between PLK1 and CUL3 (Fig 9B) and results in markedly decreased polyubiquitin modification of PLK1 under denaturing conditions (Fig 9C).

Cullin-RING ubiquitin ligases (CRLs) are the largest family of E3 ubiquitin ligases that regulate both proteolytic and non-proteolytic ubiquitin signals in a large variety of cellular processes (Jang *et al*, 2020; Jerabkova & Sumara, 2019). Accumulating evidence suggests that CUL3 emerges as a critical regulator of cell division by regulating critical mitotic kinases such as PLK1, AurA and AurB (Beck *et al*, 2013; Krupina *et al*, 2016; Maerki *et al*, 2009; Sumara *et al*, 2007; Courtheoux *et al*, 2016; Moghe *et al*, 2012). However, we still lack sufficient knowledge regarding the molecular identity and function of additional factors that act in concert with CUL3 to precisely define the cellular fate of mitotic substrates and subsequently cell cycle progression. It was recently proposed that both CRL substrate recruitment as well as CRL complex assembly are dependent on the coordinated actions of specific co-adaptors and inhibitors to ensure their function in time and space (Akopian *et al*, 2022). Here, we demonstrate that UBAP2L regulates the protein levels and localization of PLK1 but of no other mitotic targets of CUL3 including AurA and AurB (Fig 3 and EV3). Moreover, endogenous UBAP2L co-immunoprecipitates with PLK1, CUL3 and KLHL22, but not with AurB during mitosis (Fig 9A). Given the loss of interaction between CUL3 and PLK1 observed upon UBAP2L depletion (Fig 9B), our data indicate that UBAP2L might be important for the recognition of PLK1 by the KLHL22/CUL3 complex. This could, at least to some extent, explain the phenotype of PLK1 accumulating in kinetochores even after metaphase to anaphase transition in the absence of UBAP2L and could suggest that UBAP2L might act as a co-adaptor for CUL3 to ensure its access to PLK1. Further studies are needed to explore whether UBAP2L might decipher the versatility of the CUL3-based ubiquitin code during cell division.

However, the accumulation of PLK1 in non-kinetochore mitotic structures and its increased protein stability upon UBAP2L depletion, suggests that UBAP2L might also regulate PLK1 independently of the CUL3-based pathway via yet uncharacterized mechanisms. PLK1 is ubiquitylated by the APC/C E3 ubiquitin ligase in anaphase via its interaction with FZR1/CDH1, which provides the signal for the proteasomal-dependent degradation of PLK1 during mitotic exit (Lindon & Pines, 2004). One possibility would be that in the absence of UBAP2L the affinity of PLK1 towards CDH1 is reduced or shifted towards CDC20, therefore leading to increased PLK1 protein stability during mitotic exit. Still, we cannot exclude that the UBAP2L-driven proteolytic signals on PLK1 might involve other E3 ligases independent of the established APC/C mechanism, or that CUL3 might associate with unknown adaptors/inhibitors (Akopian *et al*, 2022) which in turn activate proteolytic ubiquitylation on PLK1. To our knowledge, such a dual regulation for PLK1 in terms of both stability and localization has only been described in one more study which addressed the role of NUMB in mitosis, a protein mostly known for its function in progenitor cell fate determination (Gulino *et al*, 2010). The authors show that NUMB depletion resulted in reduced PLK1 protein stability and in aberrant centrosome localization of PLK1 at both metaphase and anaphase, leading to disorganized γ-tubulin recruitment to centrosomes (Schmit *et al*, 2012). Our work is the first to report a unique role for UBAP2L in converging both proteolytic and non-proteolytic ubiquitin signals on PLK1 in order to ensure fidelity of mitotic progression. How exactly those two UBAP2L-dependent signaling cascades communicate with each other to precisely regulate PLK1 in time and space remains to be addressed in future studies.

### Possible consequences of aberrant PLK1 signaling

Mitotic perturbations are causally linked to aneuploidy and genomic instability (S. Pedersen *et al*, 2016). Phosphorylation and ubiquitylation pathways are tightly interconnected in mitosis and it is important to understand these links in the context of carcinogenesis. PLK1 is deregulated in human cancers and small molecule inhibitors targeting PLK1 are currently being explored for cancer treatment (Chiappa *et al*, 2022). However, preclinical success with currently available PLK1 inhibitors has not translated well into clinical success, highlighting the need for a complete understanding of upstream PLK1 regulatory mechanisms. In our view, combined therapies targeting other relevant pathways together with PLK1 may be vital to combat issues observed with monotherapy, especially resistance. In addition, research should also be directed towards understanding the mechanisms regulating localized activity of PLK1 and designing additional next generations of specific, potent PLK1 inhibitors to target cancer (Gutteridge *et al*, 2016).

We provide evidence that UBAP2L depletion results in aberrant PLK1 kinase activity and high levels in interphasic cells, which may ultimately cause genomic instability and cell death. What could be the potential consequences for cells entering the subsequent cell cycle in the presence of high PLK1 activity? PLK1 has a largely unexplored and unconventional functional territory beyond mitosis especially in processes such as DNA replication, transcription and damage checkpoint recovery (Kumar *et al*, 2017). Our study suggests that the accumulated micronuclei observed in UBAP2L depleted cells during mitotic exit is directly linked to aberrant PLK1 levels/activity in these cells (Fig 9). Micronuclei display highly heterogeneous features regarding the recruitment or retainment of replication, transcription and DNA damage response factors, ultimately being associated with chromosomal instability (Krupina *et al*, 2021). Interestingly, PLK1 has been shown to regulate RNAPIII-dependent transcription, switching from activation to repression based on its activatory status (Fairley *et al*, 2012), while a recent study implicated UBAP2L in the ubiquitylation and degradation of RNAPII through the recruitment of a Cullin-based ubiquitin complex (Herlihy *et al*, 2022). We could therefore speculate that cells with defective UBAP2L-PLK1 signaling would be more prone to unbalanced transcription which would further hijack their genome fidelity, a concept worth to be investigated in the future.

Finally, growing evidence suggests that UBAP2L is overexpressed in a variety of cancers where it displays oncogenic properties by interfering with signaling pathways that promote cancer cell proliferation, tumour vascularization, migration, invasion and metastasis (Guerber *et al*, 2022). While the oncogenic potential of UBAP2L renders it an attractive candidate for therapy, the results presented in our study linking its depletion to aberrant PLK1 activation and perturbed cell division, rather indicate that targeting UBAP2L might be a strategy that should be applied with caution. The pathway described in our study could maybe direct research efforts towards the synergistic inhibition of UBAP2L and PLK1 in specific cancer types.

## Materials and Methods

### Antibodies

Primary antibodies used in this study are the following: rabbit anti-UBAP2L (this study), rabbit monoclonal anti-PLK1 (4513, Cell Signaling Technology), mouse monoclonal anti-PLK1 (sc-17783, Santa Cruz Biotechnology), mouse monoclonal anti-Cyclin B1 (sc-245, Santa Cruz Biotechnology), rabbit monoclonal anti-Aurora A (4718, Cell Signaling Technology), mouse monoclonal anti-Flag M2 (F1804, Sigma-Aldrich), rabbit polyclonal anti-GAPDH (G9545, Sigma-Aldrich), mouse monoclonal anti-alpha-tubulin (T9026, Sigma-Aldrich), rabbit polyclonal anti-BubR1 (612502, BD Biosciences), human polyclonal anti-centromere (15-234-0001, Antibodies Incorporated), mouse monoclonal anti-ubiquitin (FK2) (ST1200, Millipore), mouse monoclonal anti-AIM-1 (Aurora B) (611083, BD Biosciences), rabbit polyclonal anti-Lamin B1 (ab16048, Abcam), rabbit polyclonal anti-phospho-PLK1 (5472, Cell Signaling Technology), goat polyclonal anti-PLK4 (NB100-894, Novus Biologicals), rabbit polyclonal anti-PLK3 (NBP2-32530, Novus Biologicals), rabbit polyclonal anti-PLK2 (GTX112022, GeneTex), rabbit polyclonal anti-Cyclin A (sc-751, Santa Cruz Biotechnology), mouse monoclonal anti-cyclin E (sc-247, Santa Cruz Biotechnology), rabbit polyclonal anti-GFP (ab290, Abcam), rabbit anti-CUL3 (From Sumara et al., 2007), rabbit polyclonal anti KLHL22 (16214-1-AP, Proteintech), mouse monoclonal anti-ubiquitin (clone P4D1) (3936, Cell Signaling Technology), mouse monoclonal anti-G3BP1 (611126, BD Biosciences), rabbit polyclonal anti-G3BP2 (A302-040A-M, Thermo Fisher Scientific), rabbit polyclonal anti-INCENP (2807, Cell Signaling Technology), rabbit polyclonal anti-pericentrin (ab4448, Abcam), rabbit polyclonal anti-gamma-tubulin (T3559, Sigma-Aldrich), rabbit polyclonal anti-PRC1 (sc-8356, Santa Cruz Biotechnology). Secondary antibodies used are the following: goat anti-mouse Alexa Fluor 488, goat anti-rabbit Alexa Fluor 488, goat anti-mouse Alexa Fluor 568, goat anti-rabbit Alexa Fluor 568, goat anti-human Alexa Fluor 568 (Thermo Fisher Scientific), goat anti-rabbit IgG-HRP conjugate and goat anti-mouse IgG-HRP conjugate (Biorad).

### Generation of stable cell lines and cell culture

HeLa WT and UBAP2L KO cell lines were generated using CRISPR/Cas9-mediated gene editing as described in (Fig EV1C). Two gRNAs targeting UBAP2L (5’-TGGCCAGACGGAATCCAATG-3’ and 5’-GTGGTGGGCCACCAAGACGG-3’) were cloned into pX330-P2A-EGFP/RFP (Zhang *et al*, 2017) through ligation using T4 ligase (New England Biolabs) using the following primers: hUBAP2L KO exon5 sgRNA-1-Fwd 5’-CACCGTGGCCAGACGGAATCCAATG-3’, hUBAP2L KO exon5 sgRNA-1-Rv 5’-AAACCATTGGATTCCGTCTGGCCAC-3’, hUBAP2L KO exon5 sgRNA-2-Fwd 5’-CACCGGTGGTGGGCCACCAAGACGG-3’, hUBAP2L KO exon5 sgRNA-2-Rv 5’-AAACCCGTCTTGGTGGCCCACCACC-3’. HeLa cells were transfected and GFP and RFP double positive cells were collected by FACS (BD FACS Aria II), cultured for 2 days and seeded with FACS into 96-well plates. Obtained UBAP2L KO single-cell clones were validated by Western blot and sequencing of PCR-amplified targeted fragment by Sanger sequencing (GATC) using the following primers: hUBAP2L KO exon5-DNA sequencing-Fwd 5’-CGAATGCATCTAGATATCGGATCCCTGCTGAGTGGAGAATGGTTA-3’, hUBAP2L KO exon5-DNA sequencing-Rv 5’-GCCTCTGCAGTCGACGGGCCCGGGAGACTGGTGGCAGTTGGTAG-3’.

For the generation of PLK1-eGFP KI cell line, HeLa Kyoto cells were transfected with Cas9-eGFP, sgRNA 5’-TCGGCCAGCAACCGTCTCA-3’ (targeting Plk1) and a repair templates (Genewiz). The repair template was designed as a fusion of 5xGly-eGFP flanked by two 500 bp arms, homologous to the genomic region around the Cas9 cutting site. 5 days after transfection eGFP positive cells were sorted and expanded for one week before a second sorting of single cells in a 96 well plates. After 2-3 weeks cells were screened by PCR.

All cultured cells were kept in culture in 5% CO2 humidified incubator at 37°C. HeLa Kyoto and derived stable cell lines (UBAP2L WT and KO, PLK1-eGFP KI) cell lines were cultured in Dulbecco’s Modified Eagle Medium (DMEM 4,5g/L Glucose) supplemented with 10% Foetal Calf Serum (FCS), 1% penicillin and 1% streptomycin. Human U-2 Osteosarcoma (U2OS) cells were cultured in DMEM supplemented with 1g/L Glucose, 10% FCS and 40μg/mL Gentamycin. DLD-1 cells were kept in culture in Roswell Park Memorial Institute (RPMI) medium without HEPES supplemented with 10% FCS and 40μg/mL Gentamycin.

### Cloning

Human UBAP2L isoform1 of 1087aa was isolated from HeLa cDNA and amplified by PCR. PCR products were cloned into pcDNA3.1 vector. UBAP2L FL, NT and CT fragments were generated using the following primers: hUBAP2L-FL-Flag-Fwd 5’-TTTGAATTCTTATGACATCGGTGGGCACTAACC-3’, hUBAP2L-FL-Flag-Rv 5’-TTTCTCGAGTCAGTTGGCCCCCCAGC-3’, hUBAP2L-NT-Flag-Fwd 5’-TTTGAATTCTTATGACATCGGTGGGCACTAACC-3’, hUBAP2L-NT-Flag-Rv 5’-TTTCTCGAGTTAAGCAGAAAACCTTCCTCCTCG-3’, hUBAP2L-CT-Flag-Fwd 5’-TTTGAATTCTTATGCAAGGAATGGGAACCTTTAACCCAGC-3’, hUBAP2L-CT-Flag-Rv 5’-TTTCTCGAGTCAGTTGGCCCCCCAGC-3’.

### Plasmid and siRNA transfections

Lipofectamine RNAiMAX (Invitrogen) was used to transfect siRNAs according to the manufacturer’s instructions. Final concentration of siRNA used varies from 20 to 40nM. Used oligonucleotides are the following: non-targeting siRNA (siCTL) (D-001210-02-05, Dharmacon), siUBAP2L (J-021220-09-0002, Dharmacon), siCUL3 (From Sumara et al., 2007), siG3BP1 (From Cirillo et al., 2020), siG3BP2 (From Cirillo et al., 2020).

Jetpei (Polyplus transfection) or X-tremeGENE9 (Roche) DNA transfection reagents were used to perform plasmid transfections.

### Cell synchronization

Cell cycle synchronization in G1/S, S or G2 phases was performed as previously described (Agote-Arán *et al*, 2021). For synchronization in mitosis, cells were incubated in the presence of 1μM Paclitaxel (Sigma), 1mM Monastrol (Euromedex) or 5μM STLC (Enzolifesciences) for 16h. For monastrol washout experiments, cells were collected and centrifuged at 1500rpm for 5min and resuspended in warm medium. This procedure was repeated three times and cells were cultured and collected at the desired timepoint post-release.

### Cell treatments

In order to inhibit translation, cells were treated with 100μg/mL CHX (Sigma) for the indicated time. To inhibit proteasomal degradation, cells were treated with 25μM MG132 (Tocris bioscience) for the indicated time. PLK1 enzymatic activity was inhibited using 10nM BI2536 (Euromedex) for 45min.

### Western Blot (WB)

Cells were collected by centrifugation at 1500rpm for 5 min at 4°C and washed three times with cold 1X PBS. Cell pellets were lysed using 1X RIPA buffer (50 mM Tris-HCl pH 7.5, 150 mM NaCl, 1% Triton X-100, 1 mM EDTA, 1 mM EGTA, 2 mM Sodium pyrophosphate, 1 mM NaVO4 (Na3O4V) and 1 mM NaF) supplemented with protease inhibitor cocktail (Roche) and incubated on ice for 30 min with periodic vortexing. After centrifugation at 14 000rpm for 30 min at 4 °C, total protein concentration was measured using Bradford assay by Bio-Rad Protein Assay kit (Bio-Rad). Samples were boiled for 10 min at 96 °C in 1X Laemmli buffer (LB) with β-Mercaptoethanol (BioRad) and resolved on pre-casted 4-12% Bis-Tris gradient gels (Thermo Scientific) before being transferred to a polyvinylidene difluoride (PVDF) membrane (Millipore) using wet transfer modules (BIO-RAD Mini-PROTEAN® Tetra System). Membranes were blocked in 5% non-fat milk powder, 5% bovine serum albumin (BSA, Millipore), or 5% non-fat milk powder mixed with 3% BSA for 1h at room temperature, followed by incubation with antibodies diluted in TBS-T-5% BSA/5% milk. All incubations with primary antibodies were performed for overnight at 4°C. Membranes were washed with TBS-T and developed using Luminata Forte Western HRP substrate (Merck Millipore).

### Immunoprecipitation (IP)

Cells were collected by centrifugation at 1500rpm for 5min at 4°C and washed twice with ice-cold 1X PBS and cell lysates for immunoprecipitation were prepared using IP buffer (25 mM Tris-HCl pH7.5, 150 mM NaCl, 1% Triton X-100, 5mM MgCl2, 2mM EDTA, 2mM PMSF and 10mM NaF) supplemented with protease inhibitors (Roche) and incubated on ice for 20 minutes. After centrifugation at 14000rpm for 30 minutes at 4°C cleared supernatant was used immediately. Lysates were equilibrated to volume and concentration.

For endogenous IPs, IgG and target specific antibodies (anti-UBAP2L or anti-PLK1) as well as protein G sepharose 4 Fast Flow beads (GE Healthcare Life Sciences) were used. Samples were incubated with the IgG and specific antibodies overnight at 4°C under rotation. Beads were blocked with 3% BSA diluted in 1X IP buffer and incubated for 4h at 4°C with rotation. Next, the IgG/specific antibodies-samples and blocked beads were incubated together to a final volume of 1 ml for 4h at 4°C under rotation. The beads were washed with IP buffer 4 to 6 times for 10 min each at 4°C under rotation. Notably, beads were pelleted by centrifugation at 1500rpm for 5 min at 4°C. The washed beads were directly eluted in 2X LB with β-Mercaptoethanol and boiled for 10 min at 96°C and samples were resolved by WB as described above.

For IP under denaturing conditions, HeLa cells were transfected with His/Biotin Ubiquitin and pEGFP-PLK1 or with His/Biotin Ubiquitin and pEGFP-N1 for 30h. Cells were treated with 50μM MG132 and lysed in Urea lysis buffer (8M Urea, 300mM NaCl, 50mM Na2HPO4, 50mM Tris-HCl and 1mM PMSF, pH8). Supernatants were cleared by centrifugation at 14000rpm for 15 minutes and incubated with GFP-Trap agarose beads (Chromotek) overnight at 4°C under rotation. Beads were washed by Urea lysis buffer, eluted in 2X LB and analyzed by WB.

### Subcellular fractionation

After appropriate cell synchronization, cells were collected by centrifugation at 1500rpm for 5min at 4°C. The cytosolic fraction was removed by incubation in hypotonic buffer (10mM HEPES pH7, 50mM NaCl, 0,3M sucrose, 0,5% Triton X-100) supplemented with protease inhibitors cocktail (Roche) for 10min on ice and centrifuged at 1500rpm for 5min at 4°C. The remaining pellet was washed with the washing buffer (10mM HEPES pH7, 50mM NaCl, 0,3M sucrose) supplemented with protease inhibitors cocktail (Roche). The soluble nuclear fraction was collected by incubation with the nuclear buffer (10mM HEPES pH7, 200mM NaCl, 1mM EDTA, 0,5% NP-40) supplemented with protease inhibitors cocktail (Roche) for 10min on ice and centrifuge at 14000rpm for 2min at 4°C. Lysates were boiled for 10 min at 96 °C in 1X LB with β-Mercaptoethanol (BioRad) and analyzed by WB.

### Immunofluorescence

Mitotic cells synchronized with paclitaxel, monastrol or STLC were collected with cell scrapers, centrifuged on Thermo Scientific Shandon Cytospin 4 Cytocentrifuge for 5 minutes at 1000 rpm and fixed with 4% PFA for 10 min at room temperature. Mitotic cells synchronized with DTBR were cultured on coverslips and collected as indicated below for interphasic cells. An adapted protocol was used for the experiments presented in Fig EV2. After the appropriate synchronization using DTBR, the cytoplasm was extracted from the cells to remove the large cytoplasmic fraction of UBAP2L by incubating the coverslips in cold 0,01% Triton X-100 for 1min 30sec. 4% PFA was immediately added to the coverslips after the pre-extraction and the standard IF protocol was followed.

Interphasic cells were cultured directly on coverslips, collected and fixed with 4% PFA for 10 min at room temperature. Cells were washed 3 times with 1X PBS and permeabilized with 0.5% NP-40 for 5min, rinsed with PBS-Triton 0.01% (PBS-T) and blocked with 3% BSA for 1h. Cells were then incubated with primary antibodies in blocking buffer for 1h at room temperature, washed 3 times with PBS-T and incubated with secondary antibodies in blocking buffer for 45 min at room temperature in the dark. After incubation, cells were washed 3 times with PBS-T and glass coverslips were added on cells (mitosis) or coverslips were mounted onto slides with Mowiol containing DAPI (Calbiochem).

### Live-imaging microscopy

For live-cell microscopy, cells were grown on 35mm glass bottom dishes with four compartments and SiR-DNA (Spirochrom) was added 1h before filming. For the experiments described in Figure 1, cells were synchronized with Double Thymidine and Release (DTBR) protocol and were filmed 8h after release using a 63x water immersion objective for a total time frame of 8h (one acquisition every 5 or 10min). The acquisition was done by the Yokogawa W1 rotating disk combined with a Leica 63x/1.0 water lens. Images were acquired in stacks of 25μm (2μm steps). For the experiments described in Figure 7, HeLa PLK1-eGFP KI cells were synchronized using DTBR protocol and were filmed 10h after release for a total time frame of 8h. Images were acquired every 10 min in stacks of 12μm range (0,5μm steps). Image analysis was performed using ImageJ software.

### Colony formation assay

Colony Formation Assay was performed as previously described (Pangou *et al*, 2021). Briefly, 500 cells were plated per well in a 6-well plate in technical triplicate for each condition and were cultures for 7 days. Colonies were washed using 1X PBS and fixed with 4% paraformaldehyde (PFA) and stained with 0,1% Crystal Violet for 30min. The number of colonies and individual area were counted using an automated pipeline generated on Fiji software. Three biological replicates were performed.

### Automatic nuclear intensity measurements

Quantification of nuclear intensity was conducted using CellProfiler as previously described (Agote-Aran *et al*, 2020). Briefly, the DAPI channel of pictures was used to delimit the area of interest (nucleus) and the intensity of the desired channel was automatically measured in this specific area. At least 200 cells from three biological replicates were measured per condition.

### Statistical analysis

At least three independent biological replicates were performed for each experiment. Graphs were made using GraphPad Prism and Adobe illustrator softwares. Schemes were created using BioRender.com. All experiments were done in a strictly double-blind manner. Image quantifications were carried out in a blinded manner. Normal distribution was assessed using Shapiro-Wilk test for each experiment. Normal sets of data were analyzed using two sample two-tailed T-test or One-way ANOVA with Dunnett’s or Sidak’s correction in case of multiple group analysis. For non-normal data, Mann-Whitney’s or Kruskal-Wallis test with Dunn’s correction tests were performed. Graphically, error bars represent Standard Deviation (SD) and for all experiments, significance stars were assigned as following: * P<0,05, ** P<0,01, *** P<0,001, **** P<0,0001, ns=non-significant.

## Supporting information

Extended View Figures EV1-5

## Acknowledgments

We are grateful to Monica Gotta and Luca Cirillo from the Department of Cell Physiology and Metabolism and University of Geneva that kindly generated and shared with us the HeLa PLK1-eGFP knock in cell line, as well as for helpful discussions on the manuscript. We thank the Imaging Center of the IGBMC (ICI) and the IGBMC core facilities for their support on this research. L.G. was supported by Labex international PhD fellowship from IGBMC and IMC-Bio graduate school. E.P. was supported by postdoctoral fellowships from the ANR-10-LABX-0030-INRT. Research in the I.S. laboratory was supported by IGBMC, CNRS, Fondation ARC pour la recherche sur le cancer (ARC), Institut National du Cancer (INCa), Ligue Nationale contre le Cancer, USIAS, Sanofi iAward Europe, PCSI and ANR. This work of the Interdisciplinary Thematic Institute IMCBio, as part of the ITI 2021-2028 program of the University of Strasbourg, CNRS and Inserm, was supported by IdEx Unistra (ANR-10-IDEX-0002), and by SFRI-STRAT’US project (ANR 20-SFRI-0012) and EUR IMCBio (ANR-17-EURE-0023) under the framework of the French Investments for the Future Program. E.P. would like to dedicate this work to the loving memory of Dr. Emmanuel Venieris.

## Author contributions

Conceptualization: E.P. and I.S., Software: E.G., Methodology: L.G. and E.P., Validation: L.G and E.P., Formal Analysis: L.G and E.P., Investigation: L.G., E.P., A.V., Y.L. and C.K., Writing-Original Draft: L.G., E.P. and I.S. Writing-Review and Editing: E.P., and I.S., Visualization: L.G., E.P., and I.S., Supervision: E.P. and I.S. Funding Acquisition: I.S.

## Expanded View Figures Legends

**Figure EV1.**

**A**. Representative microscopy images from high-content visual validation siRNA screen in HeLa cells for known and predicted human UBD proteins (Krupina et al., 2016b). ROIs are shown in the corresponding numbered panels. Scale bar, 10μm.

**B, C**. Validation of CRISPR-Cas9 mediated UBAP2L KO HeLa cell clones by WB analysis

**(B)** and Sanger-sequencing **(C)**. Proteins MW is indicated in kDa. WB is representative of three independent replicates.

**D, E**. DLD-1 cells were transfected with the indicated siRNAs for 48h and the presence of micronuclei was assessed by IF microscopy **(D)** and quantified in **(E)**. Scale bar, 5μm. Graphs represent the mean of three replicates ± standard deviation (SD) (two sample two-tailed t-test **P<0,01).

**F, G**. U2OS cells were transfected with the indicated siRNAs for 48h and the presence of micronuclei was assessed by IF microscopy **(F)** and quantified in **(G)**. Scale bar, 5μm. Graphs represent the mean of three replicates ± standard deviation (SD) (two sample two-tailed t-test *P<0,05).

**Figure EV2**

**A-E**. Representative IF pictures of HeLa cells synchronized in mitosis using DTBR after chemical pre-extraction of the cytoplasm using 0,01% of Triton X-100 for 1m30. UBAP2L localization was assessed by co-staining with indicated mitotic structures markers. ROIs are shown in the corresponding numbered panels. Scale bar, 5μm.

**Figure EV3**.

**A**. Representative IF images of control or UBAP2L-downregulated HeLa cells synchronized in mitosis using DTBR (n=3). Scale bar, 4μm.

**B-E**. Representative IF images of WT or UBAP2L KO HeLa cells synchronized in G1/S using DTB. Scale bar, 5μm. The percentage of cells expressing AurA, Cyclin B1 or AurB was quantified in **(C), (D)** and **(E)** respectively. At least 100 cells per condition was analyzed for each experiment. Graphs represent the mean of three replicates ± standard deviation (SD) (two sample two-tailed t-test ns=non-significant).

**Figure EV4**.

**A**. Schematic representation of UBAP2L protein fragments. Indicated numbers stand for aminoacids (aa).

**B**. WB analysis of G1/S synchronized (DTB) WT or UBAP2L KO HeLa cells transiently transfected with the indicated flag-tagged UBAP2L protein fragments. Proteins MW is indicated in kDa. Arrows point to the migration of each fragment. WB is representative of three independent replicates.

**C**. IF representative pictures of WT or UBAP2L KO HeLa cells transfected with the indicated flag-tagged UBAP2L protein fragments for 48h and synchronized in mitosis using MR. Scale bar, 8μm.

**D**. WB analysis of WT or UBAP2L KO HeLa cells synchronized in G1/S with DTB and transfected with the indicated flag-tagged UBAP2L constructs and control or G3BP1/2 siRNAs for 48h. Proteins MW is indicated in kDa. WB is representative of three independent replicates.

**E**. Quantification of PLK1 nuclear intensity from the experiment described in **(D)**. At least 150 cells were quantified per condition for each replicate. Each dot of the graph represents PLK1 nuclear intensity in a single nucleus. The measurements of three biological replicates are combined, red bars represent the mean (Kruskal-Wallis test with Dunn’s correction *P<0,05, ****P<0,0001, ns=non-significant).

**Figure EV5**.

**A**. Schematic representation of the screening strategy used to identify PLK1-eGFP positive clones. Forward (Fw) and Reverse (Rv) primers used are annotated. Bp stands for base pair. **B, C**. Agarose gel electrophoresis **(B)** and WB analysis **(C)** of PLK1 WT and PLK1-eGFP HeLa cells lysates. DNA fragments length is indicated in bp. Proteins MW is indicated in kDa.

**D**. Scatterplot representing the time from prophase to anaphase (seconds) in HeLa PLK1 WT and PLK1-eGFP cell lines. Error bars indicate Standard Error of the Mean. The number of analyzed cells is indicated in the graph. Statistical significance was determined using Mann-Whitney test (ns=non-significant).

**E**. Representative time frames of a 12 hours movie of HeLa PLK1-eGFP. Scale bar, 10μm. Time is indicated as hh:mm.

**F, G**. Representative IF pictures of WT or UBAP2L KO HeLa cells synchronized in mitosis with DTBR **(G)** and quantification of the relative PLK1 intensity at the outer kinetochore (A.U.) **(G)**. ROIs are shown in the corresponding numbered panels. Scale bar, 5μm. At least 50 cells were quantified per condition for each experiment and a minimum of 10 pairs of kinetochores per cell was analyzed. Each dot represents PLK1/BubR1 intensity ratio at a single pair of kinetochores. The measurements of three biological replicates are combined, red bars represent the mean (Mann-Whitney test ****P<0,0001).

**Movies EV1-3**.

Spinning disk time lapse microscopy of WT HeLa cells (EV1) and UBAP2L KO cells (EV2 and EV3) synchronized in mitosis with DTBR and analyzed by spinning disk live-video microscopy for 8 hours. SiR-DNA was used to stain DNA. Z stacks (25μm range, 2μm step) were acquired every 10 minutes and maximum intensity projection images are shown at speed 7 frames per second.

**Movies EV4-6**.

Spinning disk time lapse microscopy of HeLa cells transfected with siCTL or (EV4) or siUBAP2L (EV5 and EV6) for 48h, synchronized in mitosis with DTBR and analyzed by spinning disk live-video microscopy for 8 hours. SiR-DNA was used to stain DNA. Z-stacks (25μm range, 2μm step) were acquired every 5 minutes and maximum intensity projection images are shown at speed 7 frames per second.

**Movies EV7 and EV8**.

Spinning disk time lapse microscopy of PLK1-eGFP KI HeLa cells transfected with siCTL (EV7) or siUBAP2L (EV8) for 48h, synchronized in mitosis with DTBR and analyzed by spinning disk live-video microscopy for 8 hours. SiR-DNA was used to stain DNA. Z-stacks

(12μm range, 0,5μm step) were acquired every 10 minutes and maximum intensity projection images are shown at speed 7 frames per second.

