## Extended View Figures EV1-5 for "UBAP2L-dependent coupling of PLK1 localization and stability during mitosis"

**Affiliations :**
[1] Institut de Génétique et de Biologie Moléculaire et Cellulaire (IGBMC), Illkirch, France.
[2] Centre National de la Recherche Scientifique UMR 7104, Strasbourg, France.
[3] Institut National de la Santé et de la Recherche Médicale U964, Strasbourg, France.
[4] Université de Strasbourg, Strasbourg, France.
[5] Current address, Novartis, Basel, Switzerland

#### Revised Supplementary (Expanded View) Figures and Legends (EV1-5)

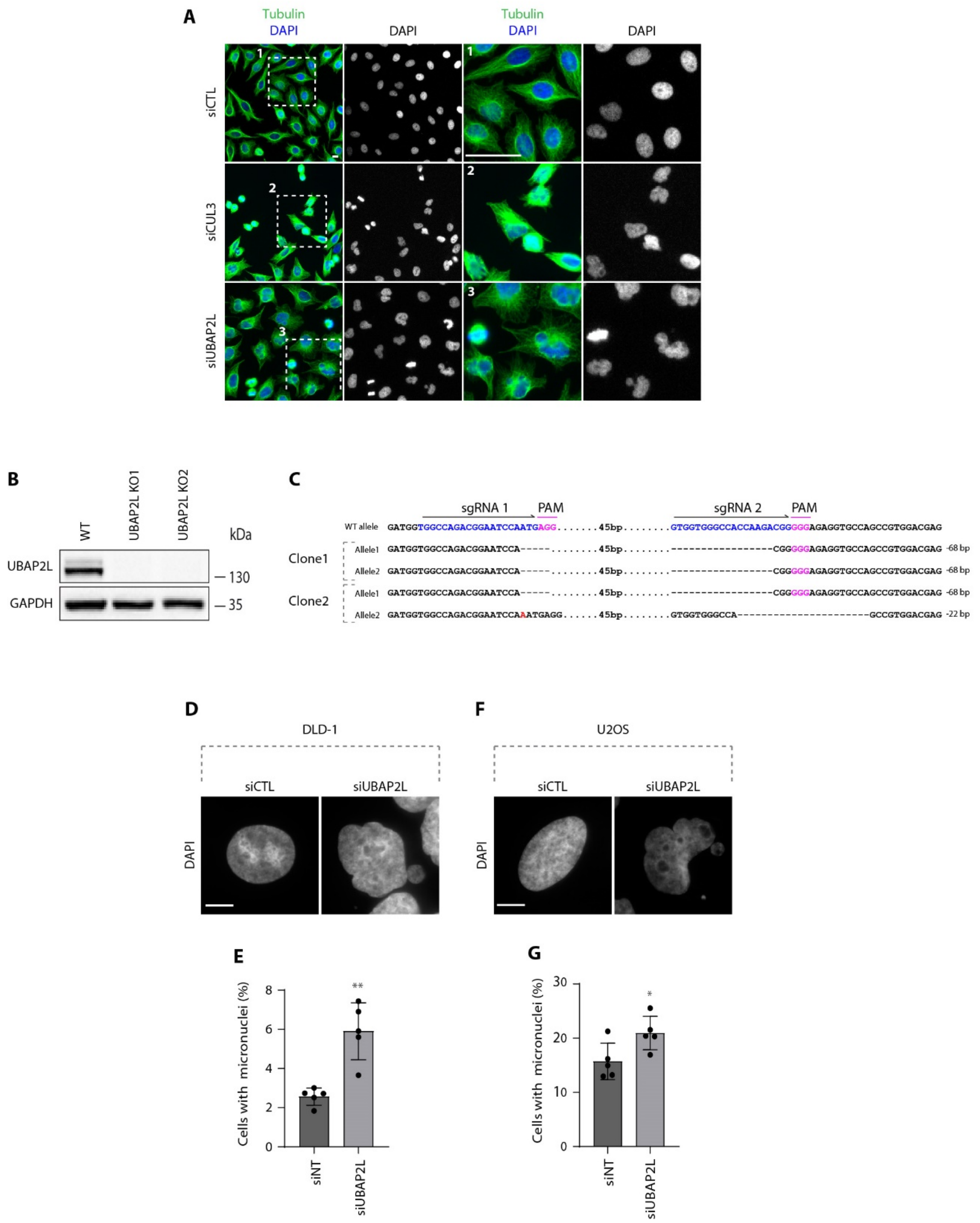


**Figure EV1.**

**A.** Representative microscopy images from high-content visual validation siRNA screen in HeLa cells for known and predicted human UBD proteins (Krupina et al., 2016b). ROIs are shown in the corresponding numbered panels. Scale bar, 10µm.

**B, C.** Validation of CRISPR-Cas9 mediated UBAP2L KO HeLa cell clones by WB analysis **(B)** and Sanger-sequencing **(C)**. Proteins MW is indicated in kDa. WB is representative of three independent replicates.

**D, E.** DLD-1 cells were transfected with the indicated siRNAs for 48h and the presence of micronuclei was assessed by IF microscopy **(D)** and quantified in **(E)**. Scale bar, 5µm. Graphs represent the mean of three replicates ± standard deviation (SD) (two sample two-tailed t-test **P<0,01).

**F, G.** U2OS cells were transfected with the indicated siRNAs for 48h and the presence of micronuclei was assessed by IF microscopy **(F)** and quantified in **(G)**. Scale bar, 5µm. Graphs represent the mean of three replicates ± standard deviation (SD) (two sample two-tailed t-test *P<0,05).

**Figure EV2.**

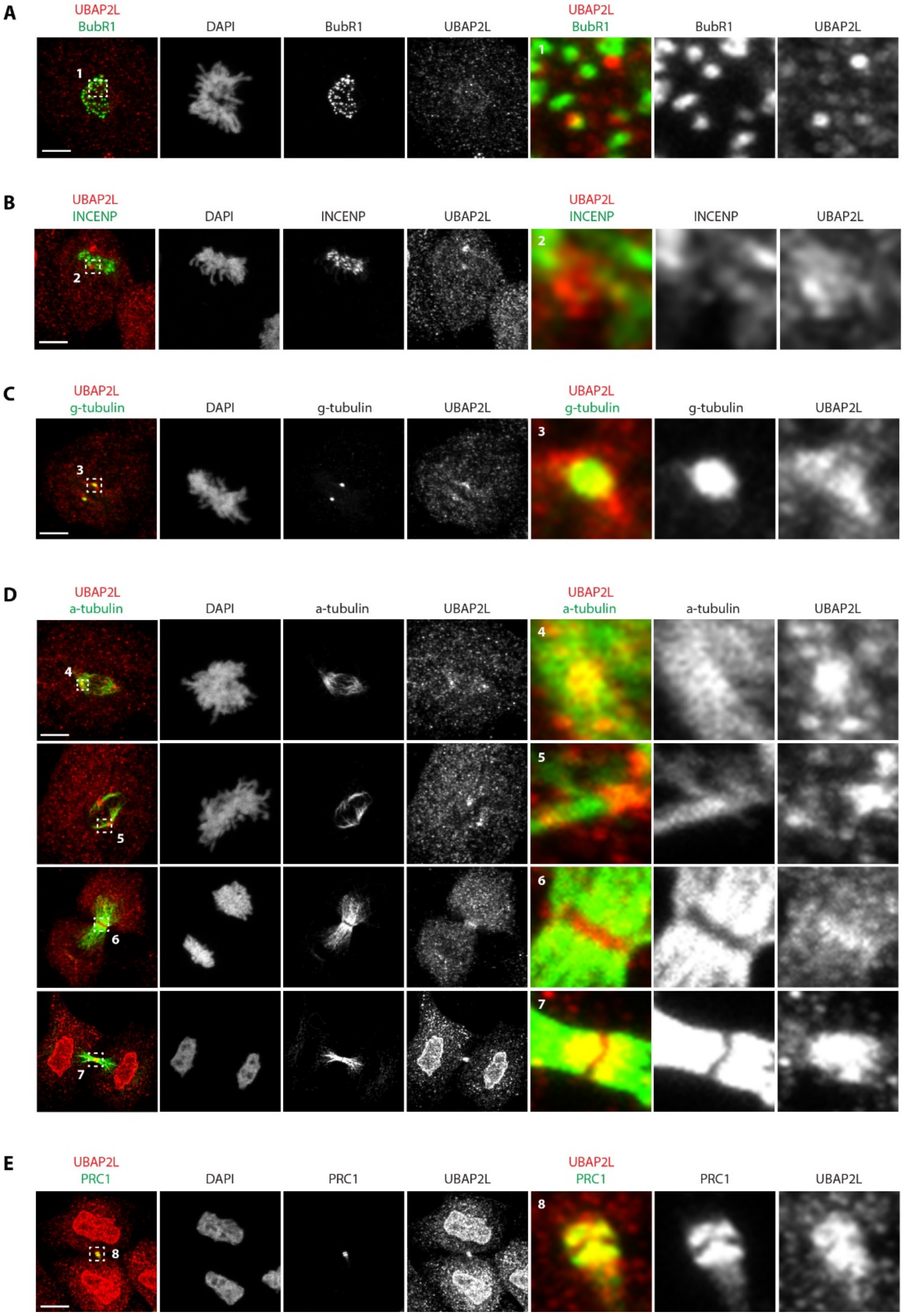


**Figure EV2.**

**A-E.** Representative IF pictures of HeLa cells synchronized in mitosis using DTBR after chemical pre-extraction of the cytoplasm using 0,01% of Triton X-100 for 1m30. UBAP2L localization was assessed by co-staining with indicated mitotic structures markers. ROIs are shown in the corresponding numbered panels. Scale bar, 5µm.

**Figure EV3.**

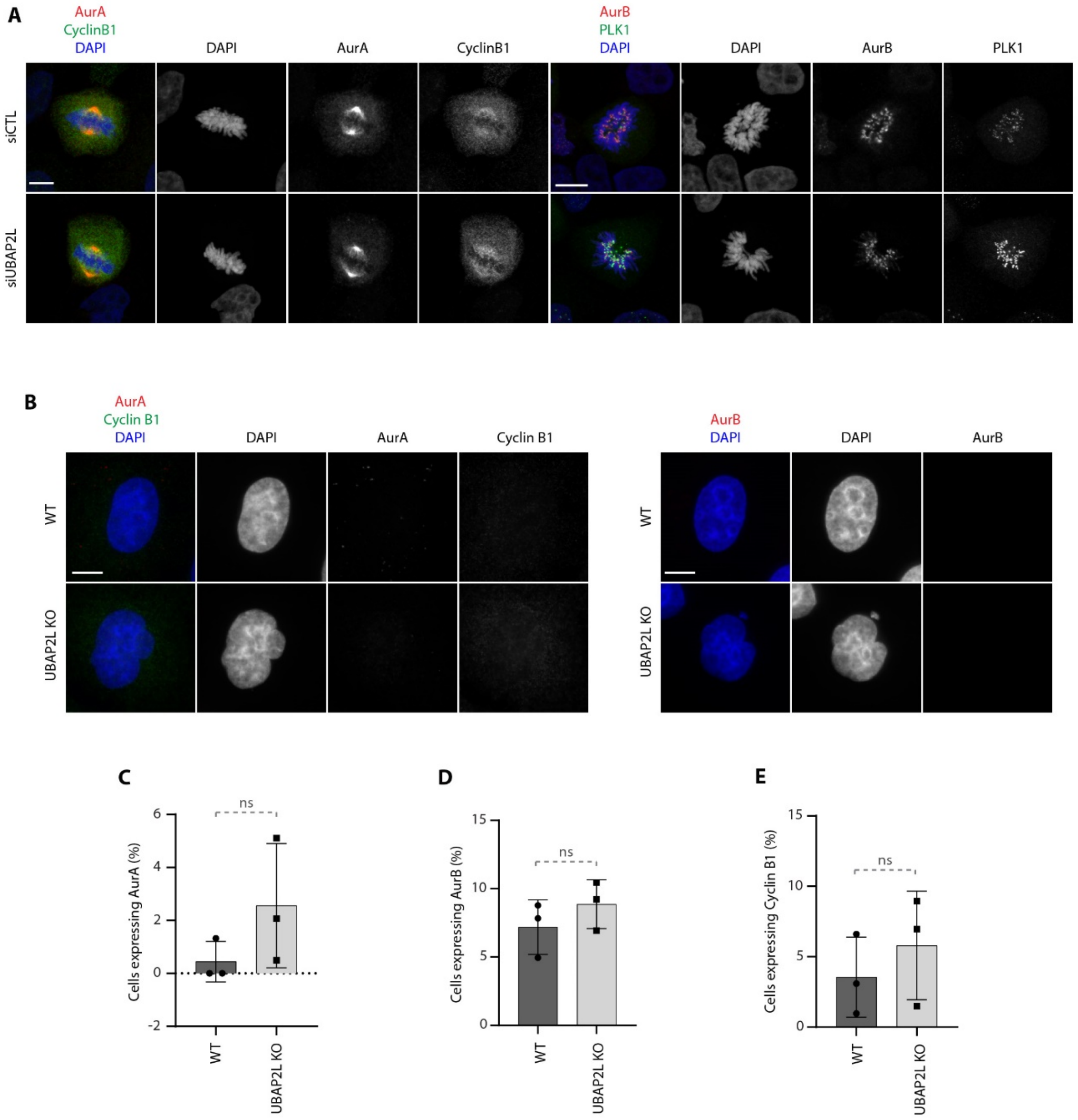


**Figure EV3.**

**A.** Representative IF images of control or UBAP2L-downregulated HeLa cells synchronized in mitosis using DTBR (n=3). Scale bar, 4µm.

**B-E.** Representative IF images of WT or UBAP2L KO HeLa cells synchronized in G1/S using DTB. Scale bar, 5µm. The percentage of cells expressing AurA, Cyclin B1 or AurB was quantified in **(C)**, **(D)** and **(E)** respectively. At least 100 cells per condition was analyzed for each experiment. Graphs represent the mean of three replicates ± standard deviation (SD) (two sample two-tailed t-test ns=non-significant).

**Figure EV4.**

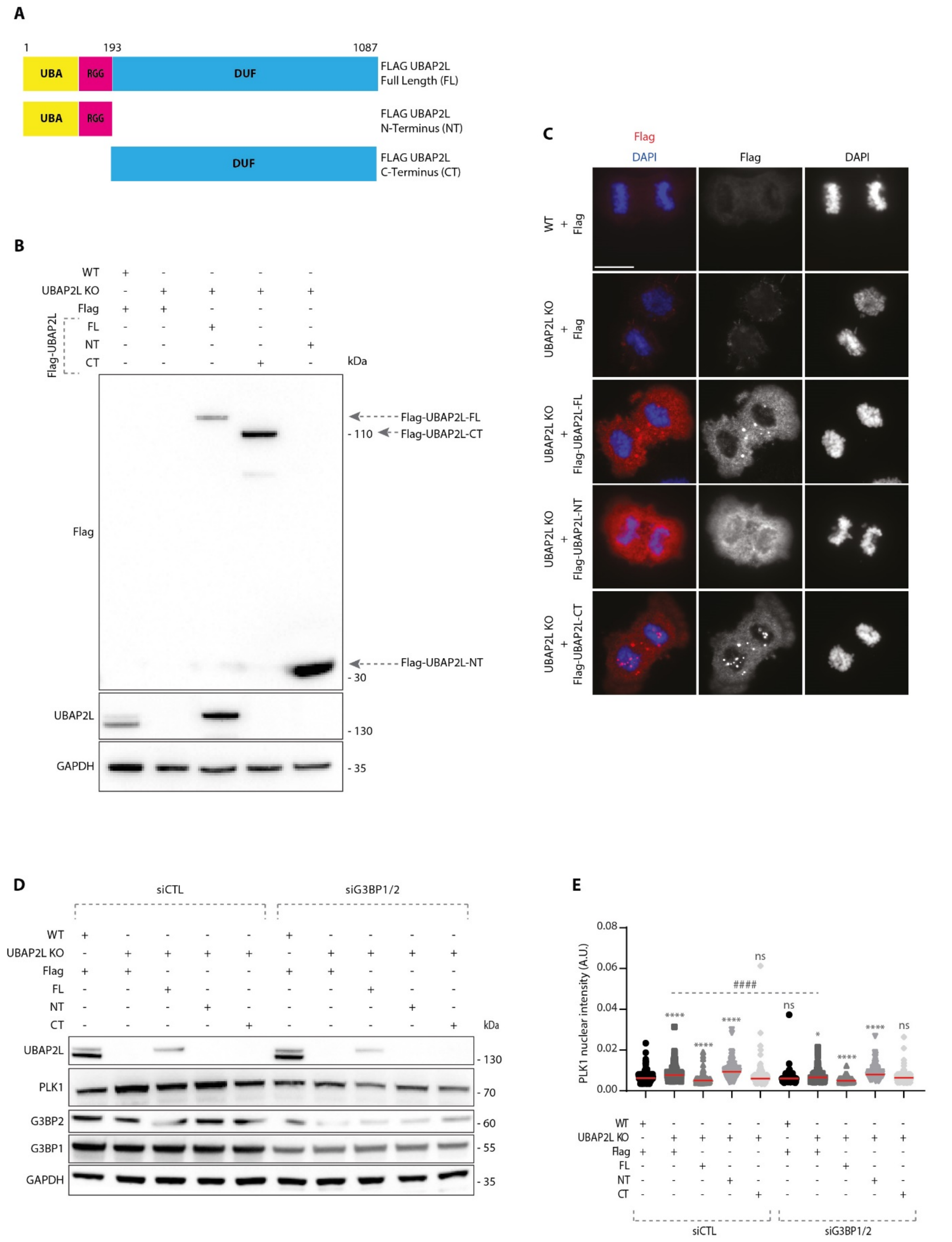


### Figure EV4.

**A.** Schematic representation of UBAP2L protein fragments. Indicated numbers stand for aminoacids (aa).

**B.** WB analysis of G1/S synchronized (DTB) WT or UBAP2L KO HeLa cells transiently transfected with the indicated flag-tagged UBAP2L protein fragments. Proteins MW is indicated in kDa. Arrows point to the migration of each fragment. WB is representative of three independent replicates.

**C.** IF representative pictures of WT or UBAP2L KO HeLa cells transfected with the indicated flag-tagged UBAP2L protein fragments for 48h and synchronized in mitosis using MR. Scale bar, 8µm.

**D.** WB analysis of WT or UBAP2L KO HeLa cells synchronized in G1/S with DTB and transfected with the indicated flag-tagged UBAP2L constructs and control or G3BP1/2 siRNAs for 48h. Proteins MW is indicated in kDa. WB is representative of three independent replicates.

**E.** Quantification of PLK1 nuclear intensity from the experiment described in **(D)**. At least 150 cells were quantified per condition for each replicate. Each dot of the graph represents PLK1 nuclear intensity in a single nucleus. The measurements of three biological replicates are combined, red bars represent the mean (Kruskal-Wallis test with Dunn's correction *P<0,05, ****P<0,0001, ns=non-significant).

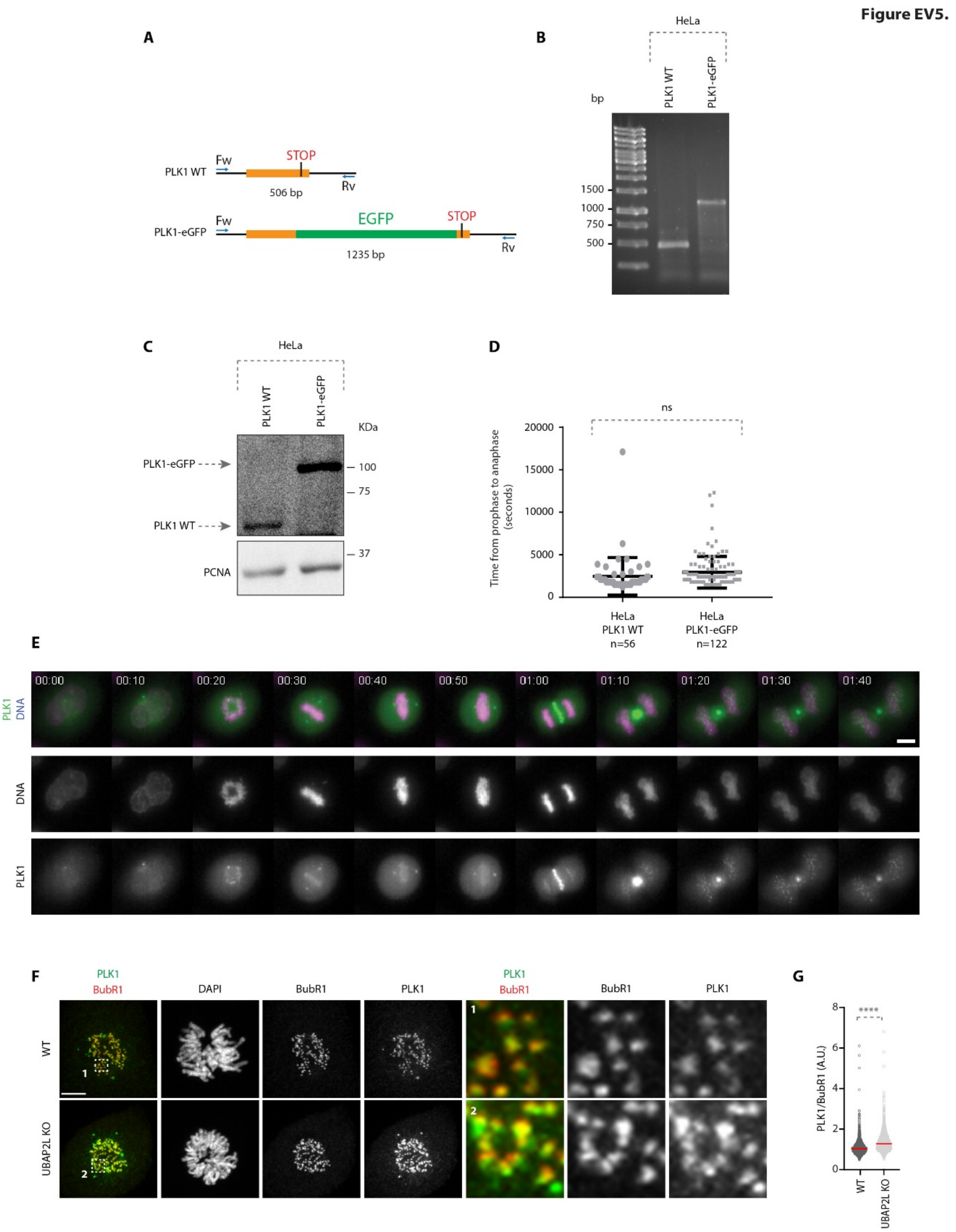


**Figure EV5.**

**A.** Schematic representation of the screening strategy used to identify PLK1-eGFP positive clones. Forward (Fw) and Reverse (Rv) primers used are annotated. Bp stands for base pair.

**B, C.** Agarose gel electrophoresis **(B)** and WB analysis **(C)** of PLK1 WT and PLK1-eGFP HeLa cells lysates. DNA fragments length is indicated in bp. Proteins MW is indicated in kDa.

**D.** Scatterplot representing the time from prophase to anaphase (seconds) in HeLa PLK1 WT and PLK1-eGFP cell lines. Error bars indicate Standard Error of the Mean. The number of analyzed cells is indicated in the graph. Statistical significance was determined using Mann-Whitney test (ns=non-significant).

**E.** Representative time frames of a 12 hours movie of HeLa PLK1-eGFP. Scale bar, 10µm. Time is indicated as hh:mm.

**F, G.** Representative IF pictures of WT or UBAP2L KO HeLa cells synchronized in mitosis with DTBR **(G)** and quantification of the relative PLK1 intensity at the outer kinetochore (A.U.) **(G)**. ROIs are shown in the corresponding numbered panels. Scale bar, 5µm. At least 50 cells were quantified per condition for each experiment and a minimum of 10 pairs of kinetochores per cell was analyzed. Each dot represents PLK1/BubR1 intensity ratio at a single pair of kinetochores. The measurements of three biological replicates are combined, red bars represent the mean (Mann-Whitney test ****P<0,0001).

**Movies EV1-3.**

Spinning disk time lapse microscopy of WT HeLa cells (EV1) and UBAP2L KO cells (EV2 and EV3) synchronized in mitosis with DTBR and analyzed by spinning disk live-video microscopy for 8 hours. SiR-DNA was used to stain DNA. Z stacks (25µm range, 2µm step) were acquired every 10 minutes and maximum intensity projection images are shown at speed 7 frames per second.

**Movies EV4-6.**

Spinning disk time lapse microscopy of HeLa cells transfected with siCTL or **(**EV4) or siUBAP2L (EV5 and EV6) for 48h, synchronized in mitosis with DTBR and analyzed by spinning disk live-video microscopy for 8 hours. SiR-DNA was used to stain DNA. Z-stacks (25µm range, 2µm step) were acquired every 5 minutes and maximum intensity projection images are shown at speed 7 frames per second.

**Movies EV7 and EV8.**

Spinning disk time lapse microscopy of PLK1-eGFP KI HeLa cells transfected with siCTL (EV7) or siUBAP2L (EV8) for 48h, synchronized in mitosis with DTBR and analyzed by spinning disk live-video microscopy for 8 hours. SiR-DNA was used to stain DNA. Z-stacks (12µm range, 0,5µm step) were acquired every 10 minutes and maximum intensity projection images are shown at speed 7 frames per second.
